# Seasonal dynamics in lettuce phyllosphere microbiota and potential transmission to the human gut

**DOI:** 10.64898/2026.03.05.709721

**Authors:** Su-A In, Ji-Woo Park, Yeo-Eun Yun, Ye-Eun Lee, Eun-Jin Park, Min-Soo Kim

## Abstract

The surface of fresh vegetable leaves harbors diverse microorganisms with the potential to influence human health through the microbiome-food-gut axis. We investigated the ecology of the bacterial and fungal microbiota on green and red lettuces (n=143) for a 12-month period using high-throughput amplicon sequencing, and assessed the potential transfer of these microbiota to the human gut. Lettuce-associated fungal and bacterial microbiota exhibited substantial temporal variation, converging into two distinct seasonal cycles: early-season (S1) and late-season (S2). Seasonal progression from S1 to S2 increased species richness in season-specific fungal and bacterial taxa, while inducing abundance shifts in persistent fungal taxa and compositional shifts in persistent bacterial taxa. These seasonal dynamics resulted in more complex and stable microbial networks, in which potentially pathogenic fungi were less frequently enriched. Comparative analyses with gut microbiota datasets from 2,831 (fungal) and 3,254 (bacterial) individuals revealed that lettuce-associated fungi and bacteria were widely detected in the human gut, with bacteria detected more frequently than fungi. Season-specific taxa were detected more frequently than persistent taxa, and microbial assembly in the gut was shaped by both neutral and deterministic processes. Notably, lettuce-associated bacteria predominantly co-occurred with non-plant glycan-degrading commensal bacteria in a season-dependent manner, and enrichment of these co-occurring taxa correlated positively with gut microbiota richness and composition. Our findings provide insights into the ecological linkages between fresh vegetable microbiota and the human gut microbiota through the food-gut axis.

## Introduction

Modern westernized lifestyles are associated with reduced human gut microbiome diversity [1, 2], partly because of diminished contact with environmental microorganisms [3, 4]. While dietary plant fibers influence gut microbiome diversity and activity by providing fermentable substrates for commensal bacteria [5], plant-based food consumption also introduces live microorganisms that may contribute [6]. However, research has focused primarily on fiber content, largely overlooking the microorganisms ingested along with these foods. Therefore, the ecological contribution of food-associated microbiomes to the human gut microbiome should be considered when defining a healthy gut microbiome [7, 8].

The phyllosphere of fresh vegetables harbors diverse microbial populations [9] shaped by environmental factors from cultivation [9, 10] through postharvest processing [10, 11]. Season is a key driver of phyllosphere microbiome assembly in field-grown plants [12], as seasonal shifts in biotic and abiotic factors act as distinct ecological filters that reshape microbiota composition prior to harvest [13]. However, whether these seasonal variations persist through postharvest processing and remain detectable in the postharvest vegetable microbiome is poorly understood [14, 15]. Fresh vegetables are consumed raw, and vegetable-associated microorganisms have recently been detected in the human fecal microbiome, with significant associations with fresh vegetable consumption [6, 16]. It is therefore plausible that the human gut is exposed to seasonally variable microbial components from the phyllosphere, and that vegetable-derived microorganisms may influence gut microbiota diversity and composition through direct ecological interactions with resident gut bacteria independent of nutrient substrate effects. Furthermore, fungal transmission from vegetables to the human gut remains largely unexplored.

Here, we characterized the fungal and bacterial microbiota of the phyllosphere of green and red lettuce collected monthly for a one-year period using high-throughput sequencing of the ITS2 region and 16S rRNA gene, and identified the effects of seasonal cycling on the diversity, assembly, and ecological functions of lettuce-associated microbiota. We further compared the lettuce-associated microbiota with human gut microbiota from 13 studies (2,831 individuals for fungal and 3,254 individuals for bacterial amplicon datasets), providing evidence for the transfer of lettuce-associated bacteria and fungi to the human gut, as well as for seasonally distinct ecological interactions with commensal gut bacteria and their contribution to gut microbiota diversity. Our findings expand current understanding of phyllosphere microbiome ecology in fresh vegetables and underscore that fresh vegetables function not only as a source of dietary fiber but also as a vector for environmental microbes to which the human gut is routinely exposed.

## Materials and Methods

### Sample collection and preparation

Fresh green (n = 72) and red (n = 71) lettuce (*Lactuca sativa*) samples were purchased monthly from six different retail stores in Daejeon, South Korea, from October 2021 to September 2022 **(Table S1)**. The samples were stored at 4 °C and processed within three days. Approximately 10 g of lettuce leaves, excluding the stem region, was aseptically excised using a sterile knife. The sample was mixed with 35 ml of Tris-EDTA buffer (Bioneer, Korea) containing 2% Tween 80 (Sigma-Aldrich, USA) and inverted thoroughly for 5 min by hand [17]. The mixture was sonicated for 10 min at maximum power (Ulmasonic Select 100 Ultrasonic Cleanser, Elma, Switzerland). Microbial cells were collected by centrifugation at 10,000 ×g at 4 °C for 10 min.

### DNA extraction, amplicon sequencing, and sequence analysis

Total DNA was extracted from the microbial pellets using a DNeasy PowerSoil Pro Kit (Qiagen, Germany), following the manufacturer’s instructions. Amplicons of the ITS2 region, V5-V6 region, and V3-V4 region of the 16S rRNA gene were prepared, following previously described [11, 20]. Sequencing was performed on an Illumina MiSeq (2 × 300 bp) for ITS2 and V5–V6 amplicons, and a NovaSeq (2 × 250 bp) for V3–V4 amplicons. Amplicon sequences were preprocessed to generate amplicon sequence variants (ASVs) using QIIME2 [21], following previously described [11, 20]. Detailed procedures for amplicon sequencing, sequence analysis, and co-occurrence network analysis are provided in Supplementary Materials and Methods.

### Species-level associations between lettuce- and gut-associated ASVs

Gut-associated fungal (n = 2,831) and bacterial (n = 3,254) amplicon datasets were obtained from 13 human cohort studies through the NCBI Sequence Read Archive **(Table S2)**. To ensure compatibility for operational taxonomic unit (OTU) clustering, a total of 77 lettuce samples representing the two seasonal clusters (S1 and S2) were selected, and their V3-V4 regions were sequenced again. Read processing was conducted in the same manner as described above. Bacterial and fungal ASV sequences from the lettuce and human gut datasets were co-clustered with reference sequences from the SILVA 132 (release 2020-11-02) or UNITE 8.3 (release 2021-05-10) databases into OTUs at 97% identity using *pick_open_reference_otus.py* in QIIME1. OTUs were retained for downstream analyses only if they met the following criteria: (1) they consisted of fewer than three ASVs, and (2) the lettuce-associated ASVs within OTUs were assigned to the same seasonal cluster. Gut-associated ASVs that were co-clustered with lettuce-associated ASVs into the same OTUs were regarded as potential lettuce-derived ASVs. The V4 region sequences of the 16S rRNA gene from lettuce-associated and paired gut-associated ASVs were used for phylogenetic tree construction using the *align-to-mafft-fasttree* action in QIIME2, which employs the MAFFT algorithm for multiple sequence alignment and the FastTree algorithm for tree construction. Visualization of the phylogenetic tree was performed using iTOL (https://itol.embl.de/). The environmental sources of bacterial and fungal genera were determined based on the Global Catalogue of Microorganisms (GCM) [18]. Bacterial genera capable of plant-derived glycan degradation were manually defined based on published studies supported by experimental evidence (**Table S3**).

### Statistical analysis

*P*-values < 0.05 were considered significant. Two-tailed Mann–Whitney *U* test, Kruskal–Wallis tests with Dunn’s post-hoc tests, and one-way ANOVA with Tukey’s post-hoc test were performed using GraphPad Prism version 5.0 for Windows (GraphPad Software, SD, USA). Principal coordinate analysis (PCoA) was performed using the R package ‘vegan’. Statistical significance for distance matrices was assessed using ‘vegan::adonis’, and significance for pairwise comparisons was assessed using ‘pairwiseAdonis::pairwise.adonis’. A *z*-score was used to standardize the meteorological parameters. Procrustes analysis between ordinations was performed using ‘vegan::protest’. Distance-based redundancy analysis (db-RDA) was performed using ‘vegan::capscale’. The best-fit model was calculated using forward selection with ‘vegan::ordiR2step’. Classification into either ‘season-specific’ or ‘persistent’ categories was determined using chi-square or Fisher’s exact tests based on frequency values and MaAsLin2 [19] analysis based on log-transformed enrichment values. The log-transformed abundances of bacterial families were correlated with the CAP1 axis of a db-RDA ordination constrained to season using ‘stats::cor’, and permutation tests were conducted using ‘vegan::envfit’. For the comparisons of gut microbiota compositions between samples containing and lacking lettuce-associated OTUs, lettuce-associated OTUs were excluded, and equal sample sizes from both groups were obtained via 10 iterations of random subsampling using the ‘replicate’ function in R. The detailed procedures for logistic regression for season-associated ASV detection are described in the Supplementary Materials and Methods.

## Results

### The fungal and bacterial microbiota of green and red lettuce are abundant, host-specific, and temporally variable

We collected 143 green and red lettuce samples from six grocery stores monthly for one year (Fig. S1A and Table S1). Diverse fungal and bacterial colonies were observed (Fig. S1B), with large numbers of fungal (average ± s.d., 3.4 ± 0.8 log CFUs and 8.0 ± 1.4 log genomes) and bacterial (5.0 ± 1.0 log CFUs and 7.7 ± 1.4 log genomes) cells detected per gram of leaf tissue (Fig. S1C and D), suggesting that lettuce leaves represent favorable environments for microbial colonization. In total, 5,016 fungal and 5,610 bacterial ASVs (173 ± 116 and 173 ± 85 ASVs per sample) were obtained through high-throughput sequencing of the ITS2 region and 16S rRNA gene, respectively (Table S4). Species accumulation curves approached a plateau as the sample size increased (Fig. S1E), indicating that fungal and bacterial populations ranging from abundant to rare were sufficiently captured. The fungal and bacterial microbiota of lettuce clustered separately from those of 14 fruits and vegetables at the genus-level (*P* = 0.001) [20, 21] (Fig. S2A and Table S5). This distinctiveness was more pronounced when compared with fresh vegetables alone (*P* = 0.001) (Fig. S2B). The fungal genera *Cladosporium* and *Bulleromyces* were relatively abundant in lettuce (Fig. S2C), and the bacterial genera *Masillia* and *Sphingomonas* were uniquely present (Fig. S2D), indicating that lettuce hosts distinct fungal and bacterial communities that may reflect host-specific adaptation.

Compositional variations in the fungal and bacterial microbiota were assessed against host-associated and environmental factors using redundancy analysis. For the fungal microbiota, the best-fit model partitioned community variation along the first db-RDA axis constrained by sampling month (R^2^ = 0.3206), and the second axis constrained by distributor (R^2^ = 0.0580) (*P* = 0.001) (Fig. S2E). For the bacterial microbiota, the best-fit model partitioned community variation along the first db-RDA axis constrained by sampling month (R^2^ = 0.2185), followed by the second axis constrained by distributor (R^2^ = 0.0671), and the third axis constrained by vegetable type (R^2^ = 0.0114) (*P* = 0.001) (Fig. S2F). These results demonstrate that the lettuce fungal and bacterial microbiota exhibit pronounced temporal variation over the course of a year.

### Seasonal cycling in the lettuce fungal and bacterial microbiota

The temporal variation in the green and red lettuce microbiota largely followed a cyclical pattern, converging rather than diverging over time (Fig. 1A). Specifically, *k*-medoids clustering revealed two optimal clusters for both microbiota [22] (Fig. 1B); most samples within these clusters (87.3% of green lettuce, 75.6% of red lettuce; Table S6) corresponded to their respective sampling periods: early-season (December to June, S1) and late-season (July to November, S2). This seasonal cycling was also evident in ecological guild-based compositions (Fig. S3), with the majority of samples aligning with early- and late-season clusters (Fig. 1C and Table S7). These patterns were reflected in the diversity indices, driven primarily by changes in ASV richness (Fig. S4A and B). The total microbial abundance also exhibited a seasonal pattern, though less consistent than the compositional variation (Fig. S4C). These results suggest that temporal variation in the lettuce microbiota follows a seasonal cycle throughout the year.

**Figure 1.**
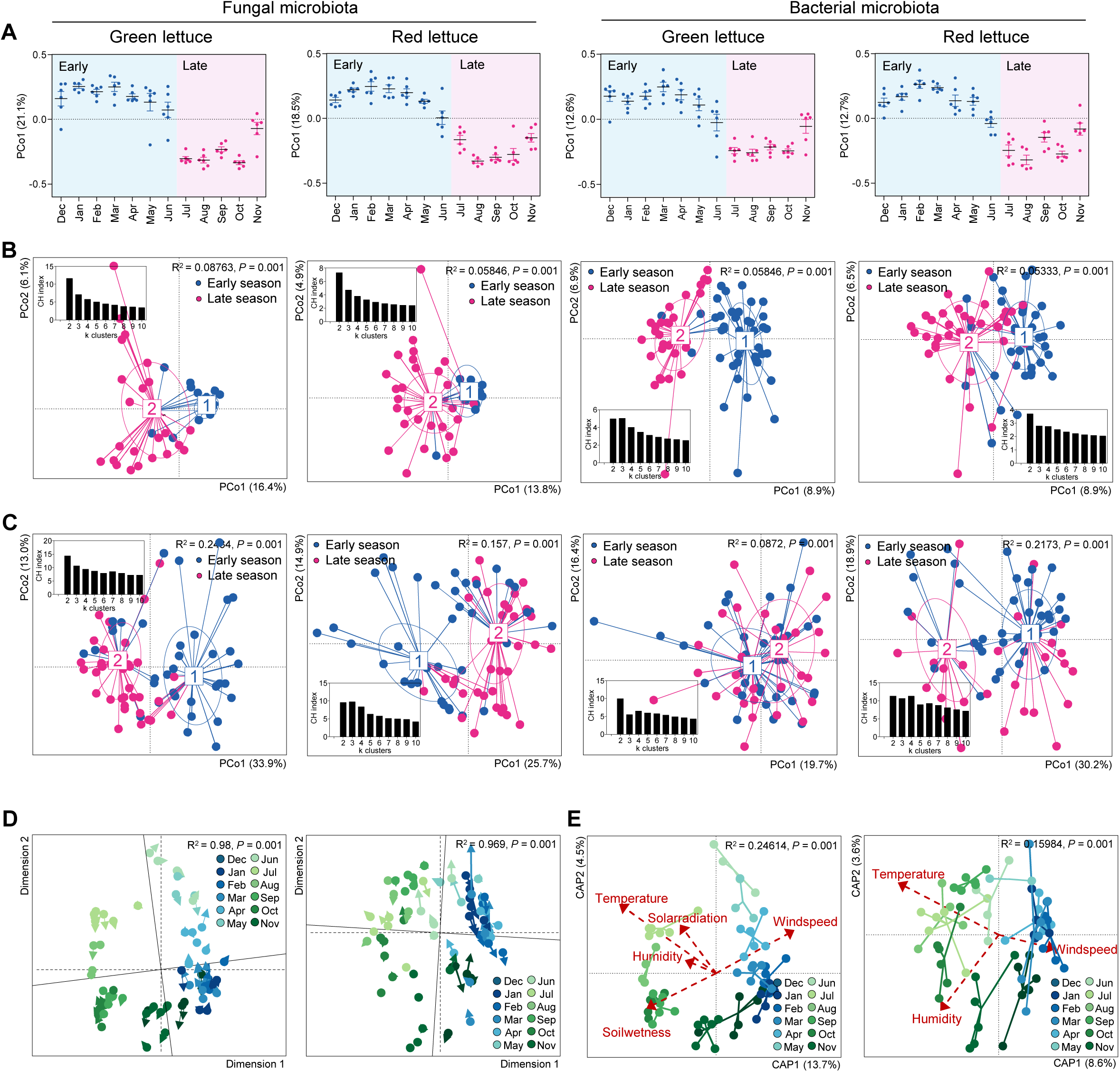
Fungal and bacterial microbiota undergo seasonal cycling in the lettuce phyllosphere. **(A)** Principal Coordinate Analysis (PCoA) plots of fungal and bacterial microbiota collected for a 12-month period, illustrating sample distribution along the first principal coordinate (PCo1). **(B)** Partition Around Medoids (PAM) clustering of fungal and bacterial microbiota and **(C)** corresponding ecological guild composition based on the k-medoids algorithm. Inset plots show the Calinski-Harabasz (CH) index used to select the optimal number of clusters (*k*). **(D)** Procrustes analysis comparing ordinations constrained by agrometeorological parameters and by sampling month for fungal and bacterial microbiota. **(E)** Redundancy analysis of fungal and bacterial microbiota constrained by agrometeorological factors. Data are presented as mean ± s.d. Statistical significance was determined by PERMANOVA and Procrustes correlation.

Given that lettuce has a relatively short production cycle from farm to consumer [23], our monthly collected samples are expected to reflect their preharvest microbiome. We examined ecological associations between the lettuce microbiota and local agrometeorological conditions by analyzing five parameters (temperature, relative humidity, wind speed, solar radiation, and soil wetness) corresponding to sample collection dates (Fig. S5 and Table S8). Procrustes analysis comparing two db-RDA models, one fitted on the agrometeorological parameters and one on the sampling month, showed strong correlations for both microbiota (R^2^ > 0.9, *P* = 0.001) (Fig. 1D). Additionally, five parameters for the fungal microbiota and three parameters for the bacterial microbiota were significantly correlated with db-RDA ordinations constrained by sampling month (*P* < 0.01) (Fig. 1E). These results support our hypothesis that the seasonal patterns observed in the lettuce microbiota may reflect season-dependent preharvest microbiota persisting through the postharvest stage.

### Seasonal enrichment patterns differ between fungal and bacterial microbiota in lettuce

Prevalence-based comparison revealed that the lettuce microbiota contained fungal and bacterial ASVs that were either season-specific or shared between S1 and S2 (referred to as “persistent”) (*P* < 0.05) (Fig. 2A). Approximately three-quarters of the ASVs were season-specific, with two-thirds of these being more prevalent in S2, indicating a seasonally increasing diversity. In contrast, one-quarter of the ASVs were persistent, yet their composition differed significantly between S1 and S2 (*P* = 0.001). Specifically, the compositional shift of the persistent fungal ASVs was driven by increased species richness and abundance from S1 to S2 (*P* < 0.05) (Fig. 2B). These patterns were evident in prevalence–density distribution plots, wherein the persistent fungal ASVs exhibited a bimodal distribution skewed towards S2 (*P* < 0.05) (Fig. 2C), and were reflected in the microbial diversity driven by fungal ASV richness (Fig. S6A). No such shifts were observed in the persistent bacterial ASVs (Fig. 2B and C); however, *Sphingomonaceae* and *Enterobacteriaceae* were commonly enriched from S1 to S2 (Fig. S6B).

**Figure 2.**
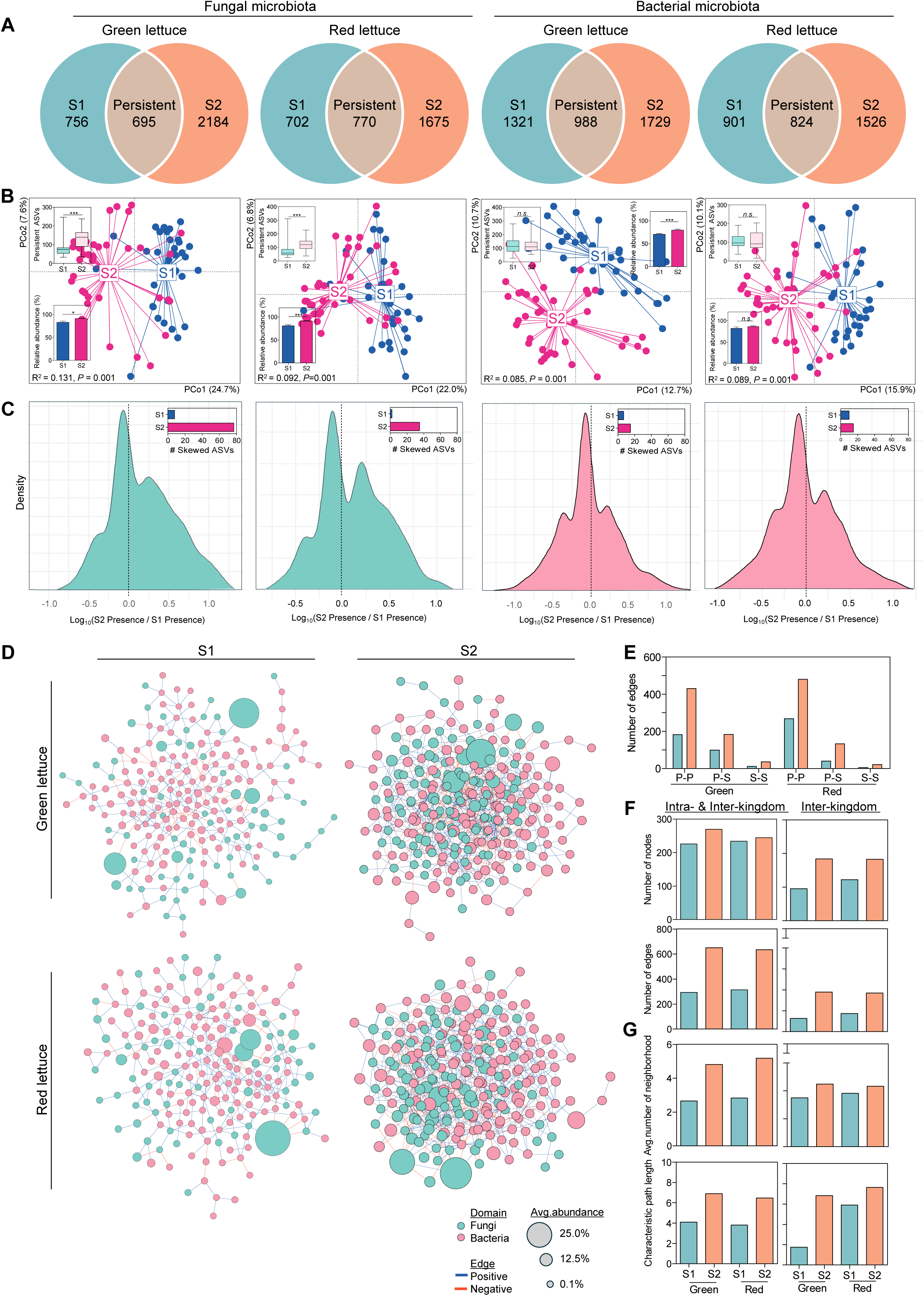
Seasonal dynamics of fungal and bacterial microbiota in the lettuce phyllosphere. **(A)** Venn diagrams showing season-specific (S1, S2) and persistent ASVs in fungal and bacterial microbiota. **(B)** PCoA plots of fungal and bacterial persistent microbiota according to season. Boxplots and barplots show comparisons of richness and total relative abundance, respectively. **(C)** Density plots of log-transformed presence ratios (S2/S1) of persistent ASVs in fungal and bacterial microbiota. Boxplots indicate the number of abundance-skewed ASVs in each season. **(D)** Co-occurrence networks of the S1 and S2 microbiota. Node size is proportional to ASV relative abundance. **(E)** Number of edges for the S1 and S2, categorized as persistent-persistent (P-P), persistent-season-specific (P-S), or between-season-specific (S-S) associations. (**F)** Comparison of the number of nodes (upper panels) and edges (lower panels), separated by all associations (left panels) and inter-kingdom associations only (right panels). **(G)** Average number of neighbors and characteristic path length for all associations and inter-kingdom associations only. Data are presented as mean ± s.d. Statistical significance was determined by Chi-square test, PERMANOVA, and two-tailed Mann-Whitney *U* tests. Symbol: n.s., not significant; *, *P* < 0.05; **, *P* < 0.01; and ***, *P* < 0.001.

Seasonal shifts in the persistent ASVs were also associated with the microbiota assembly process. In the fungal microbiota, the goodness-of-fit (R^2^) values of the Sloan neutral model [24] decreased from S1 (R^2^ = 0.582 for green lettuce and R^2^ = 0.522 for red lettuce) to S2 (R^2^ = 0.468 and R^2^ = 0.406), largely because of an increased proportion of the persistent ASVs deviating above the neutral model (Fig. S7A and C). Similar patterns were observed in the bacterial microbiota, primarily driven by persistent ASVs (Fig. S7B and D). These model predictions suggest seasonal shifts toward stronger environmental and/or host-driven selection for persistent members, which may be linked to the seasonal cycling patterns characterized by enrichments in fungal diversity and bacterial composition. Furthermore, these seasonal patterns ultimately strengthened the microbial community structure. The co-occurrence networks of the S2 microbiota showed more associations among season-specific and persistent pairs (Fig. 2D and E), as well as among inter- and intra-kingdom pairs (Fig. 2F), despite containing a similar number of nodes. This resulted in higher average neighbor numbers and characteristic path length in S2 than in S1 (Fig. 2G), suggesting that seasonal shifts toward stronger environmental and/or host-driven selection may promote both inter-and intra-kingdom interactions in the fungal and bacterial microbiota.

Opportunistic *r*-strategist pathogens can invade or emerge in the phyllosphere microbiota when their diversity and interspecies interactions are largely compromised in fresh produce [25, 26]. We assessed whether the ecological stability of the fungal and bacterial microbiota changed across seasons by identifying potentially pathogenic taxa and examining their occurrence patterns. In the fungal microbiota, similar numbers of pathogenic genera were detected in S1 (12 in green and seven in red lettuce) and S2 (ten and four); however, greater numbers of pathogenic genera were enriched in S1 (11 and seven) than in S2 (only one in green) (Fig. S8A). In contrast, far fewer pathogenic genera were detected in the bacterial microbiota, which showed minimal seasonal variation in their occurrence and enrichment patterns (Fig. S8B). Overall, our findings suggest that seasonal dynamics, together with high microbial diversity, environmental and/or host-driven selection, and strong trans-kingdom interactions, may collectively suppress the emergence of opportunistic fungal pathogens in the community.

### The fungal and bacterial microbiota of lettuce are transferred to the human intestine

Given that lactic acid-producing bacteria from fermented foods can colonize the human gut [27, 28], dietary transmission of lettuce-associated microbiota to the human gut is also plausible. We collected 13 human fecal microbiota datasets of the ITS region from 2,831 individuals and of the 16S rRNA gene from 3,254 individuals, representing diverse geographic regions and ethnicities (Fig. 3A and Table S2). We generated 16S amplicon data targeting the V3-V4 regions from the same lettuce samples, and performed species-level OTU clustering by combining ASV sequences from the lettuce and fecal microbiota (see Materials and Methods). An average of 25.2 ± 13.9 lettuce-associated fungal OTUs and 18.1 ± 13.1 bacterial OTUs were identified per study, accounting for 2.0 ± 1.5% and 1.0 ± 0.9% of the total OTU abundance, respectively (Fig. 3B and C); eight fungal and 24 bacterial OTUs were shared across at least two studies. Lettuce-associated fungal and bacterial OTUs were observed in 33.1 ± 13.8% and 67.5 ± 9.4% of the human subjects, respectively, comprising 1.5 ± 5.6% and 1.39 ± 4.2% of the total abundance, consistent with values reported previously [29] (Fig. 3D and E). Accumulation curves based on subjects and studies revealed continuously increasing trajectories for lettuce-associated OTUs (Fig. 3F), suggesting that current detection rates represent only a fraction of actual transmission.

**Figure 3.**
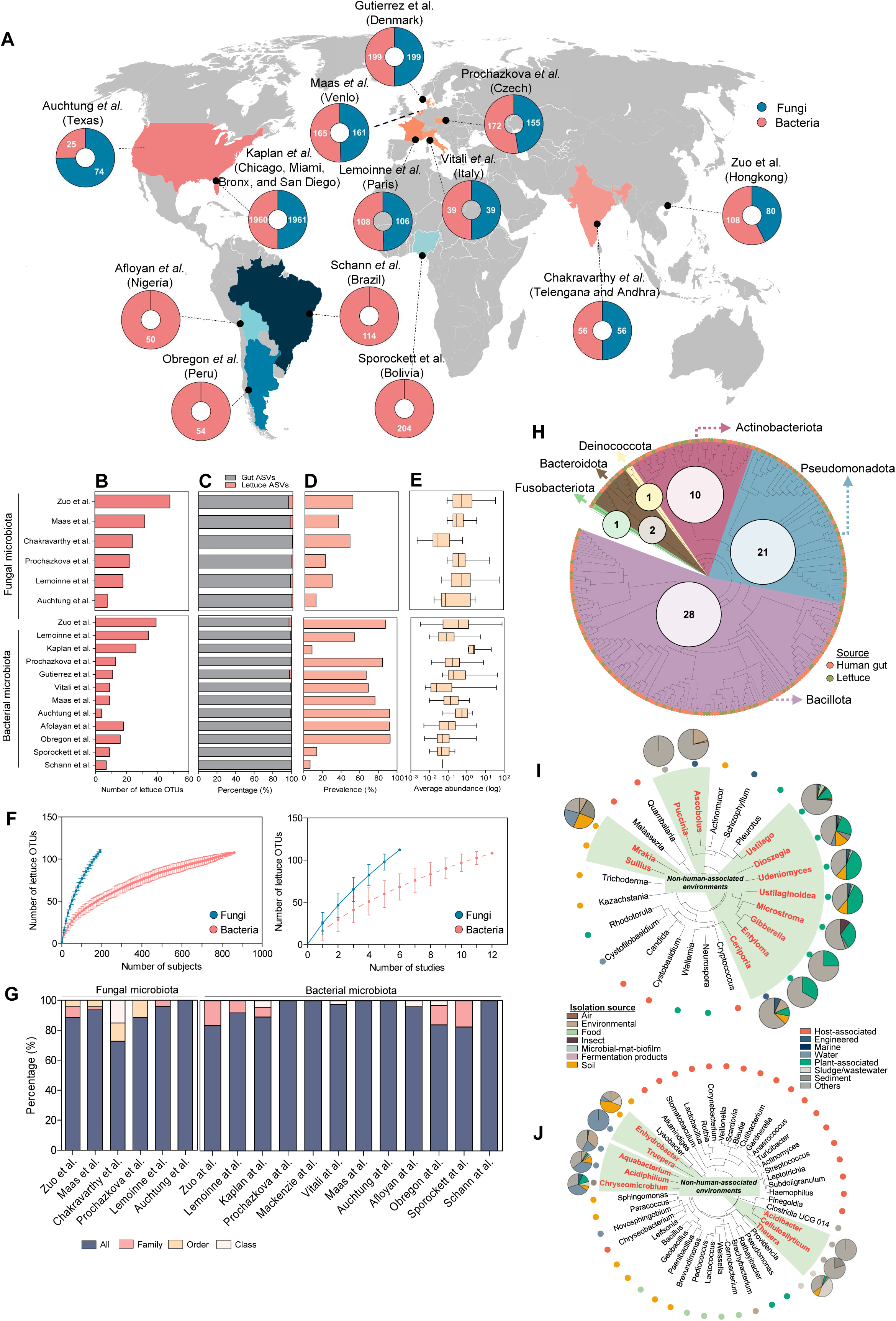
Transfer of lettuce-associated microbiota to the human gut. **(A)** Geographic distribution of human gut microbiota datasets (16S rRNA and ITS) collected from 13 studies. Pie charts indicate the number of samples in fungal and bacterial datasets. **(B)** Number of lettuce-associated fungal and bacterial OTUs identified in each study. **(C)** Proportion of lettuce-associated OTUs relative to total gut ASVs. **(D)** Prevalence of lettuce-associated OTUs across study participants. **(E)** Relative abundance of lettuce-associated OTUs in the human gut microbiota. (**F**) Accumulation curves of lettuce-associated bacterial and fungal OTUs based on the numbers of subjects and studies. **(G)** Taxonomic consistency between lettuce- and gut-associated fungal and bacterial ASVs within shared OTUs at the genus level. **(H)** Phylogenetic tree of 183 lettuce-associated bacterial OTUs shared between lettuce and the human gut. Internal circles indicate OTU counts within specific clades. Environmental isolation sources of lettuce-associated **(I)** fungal and **(J)** bacterial genera. Green shading indicates naturally occurring environment-associated clades; pie charts and colored dots show isolation source distributions based on the GCM database. Data are presented as mean ± s.d.

The taxonomic lineages of lettuce-derived ASVs were consistent with those of their gut-derived counterparts assigned to the same OTUs down to the genus-level (89.9 ± 0.03% in fungal and 91.1 ± 11.1% in bacterial OTUs) (Fig. 3G). The taxonomic relatedness of 183 bacterial OTUs was further investigated through phylogenetic analysis, revealing that 63 (34.4%) gut-associated bacterial ASVs co-localized within the same terminal node as their lettuce-associated counterparts (Fig. 3H). We additionally examined the environmental sources of bacterial and fungal genera to which lettuce-associated OTUs were taxonomically assigned on the basis of GCM [18]. Of 26 fungal genera analyzed (>1% prevalence), 12, such as *Mrakia* and *Puccinia*, have been observed in limited natural environments, particularly those exclusive to non-human-associated environments (Fig. 3I). While *Candida* spp. originate from various environments, including host-associated niches, their transmission through plant-based diet consumption has been previously reported [30]. Of 45 bacterial genera examined, eight, such as *Enhydrobacter* and *Thauera*, were exclusively of non-host-associated origins (Fig. 3J). Members of the bacterial orders *Lactobacillales*, *Eubacteriales*, *Actinomycetales*, *Bacteroidales*, and *Burkholderales* dominated the lettuce-associated OTUs, corresponding to the fruit- and vegetable-associated bacteria observed in the human gut datasets [6] (Fig. 4F). One lettuce-derived *Escherichia*-*Shigella* ASV, generally associated with leafy vegetable contamination [12], co-clustered with a gut-derived ASV, indicating minimal transfer of such bacteria in this study. Collectively, these findings suggest that closely related lettuce-associated bacterial and fungal strains may be present in the human gut microbiota, albeit at low abundance.

**Figure 4.**
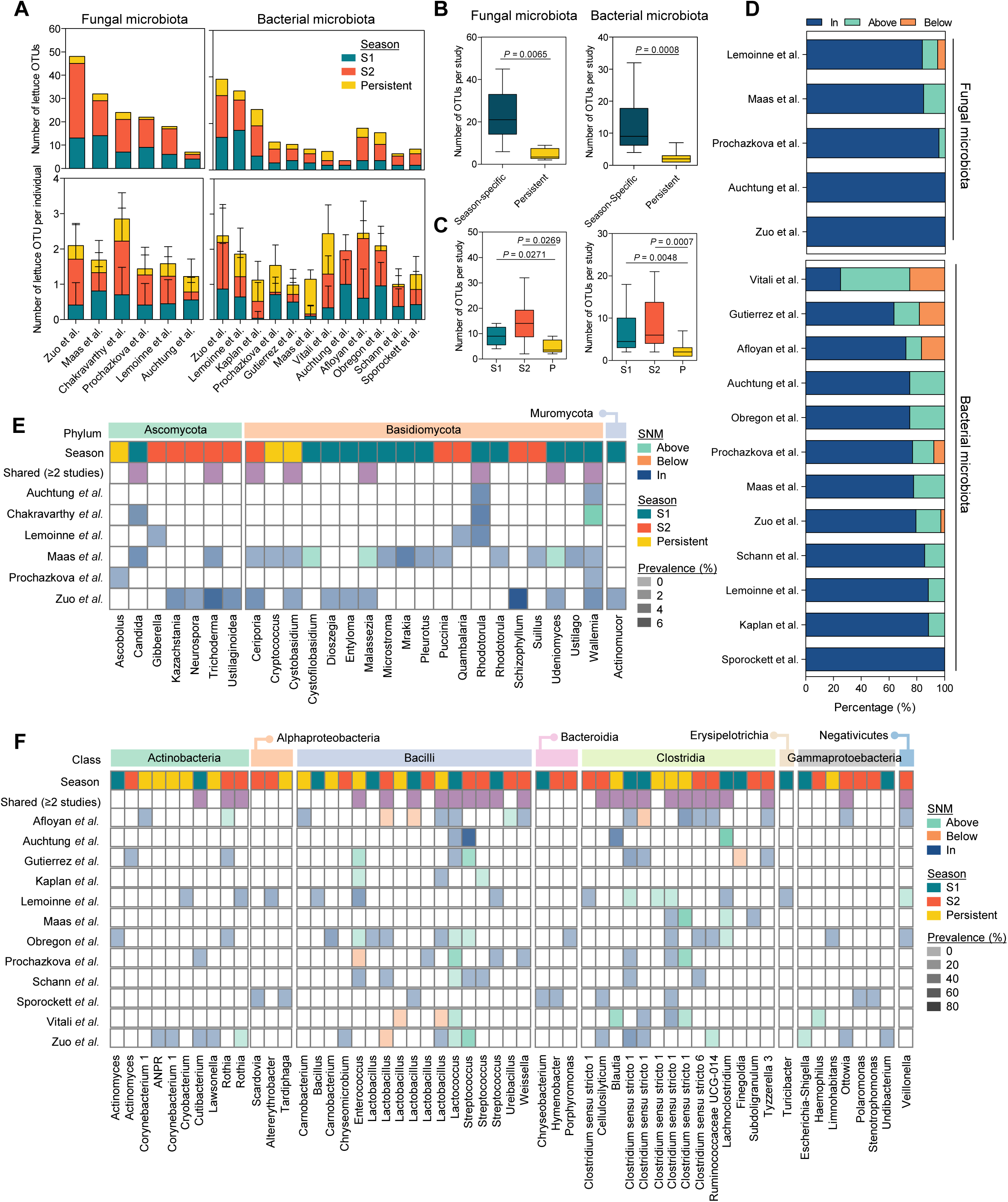
Seasonal transfer patterns of lettuce-associated fungal and bacterial OTUs into the human gut. **(A)** Numbers of season-specific (S1, S2) and persistent fungal and bacterial OTUs detected across studies (upper panel), and per individual (lower panel). **(B)** Numbers of season-specific and persistent OTUs per study. **(C)** Numbers of S1-specific, S2-specific, and persistent OTUs per study. **(D)** Proportions of fungal and bacterial OTUs relative to neutral expectations across study cohorts. Prevalence heatmaps of **(E)** fungal and **(F)** bacterial genera across cohorts based on SNM. Data are presented as mean ± s.d. Statistical significance was determined by one-tailed Mann-Whitney *U* test.

### Seasonal patterns in the transmission of lettuce-associated microbiota to the human gut and potential interactions with resident gut bacteria

We next examined whether the seasonal patterns observed in the lettuce microbiota are reflected in the human gut microbiota. The lettuce-associated OTUs were classified as either season-specific or persistent OTUs based on their corresponding lettuce-derived ASVs, resulting in both types detected across all the studies (Fig. 4A). On average, the numbers of season-specific OTUs per study and per individual significantly exceeded those of persistent OTUs (*P* < 0.05) (Fig. 4B and S9A). In particular, S2-specific OTUs were observed more frequently than S1-specific OTUs across the studies and individuals, particularly in the bacterial microbiota (Fig. 4C and S9B). This pattern was supported by the study-shared fungal (7/8, 87.5%) and bacterial OTUs (19/24, 79.2%) (Fig. 4E and F). These results reflect the seasonal increase in species richness and abundance observed in the lettuce microbiota, suggesting that microbial transmission from lettuce to the gut may occur in a season-dependent manner.

Lettuce consumption transfers both lettuce-associated microbiota and plant polysaccharides to the gut [6, 16]. While our findings indicate that microbial composition varies seasonally, polysaccharide composition is expected to remain relatively consistent. We therefore examined seasonal variation in gut-associated OTUs that co-occurred directly with lettuce-associated OTUs to distinguish microbial-driven from polysaccharide-driven influences on the gut microbiota. In total, 140 gut bacterial OTUs (33.2 ± 16.0 per study) formed co-abundance groups with 69 lettuce-associated bacterial OTUs (Fig. 5A), and 65 fungal OTUs (15.0 ± 10.7 per study) formed co-abundance groups with 47 lettuce-associated fungal OTUs (Fig. S10A), revealing more frequent co-occurrence associations in bacterial OTUs (2.48 ± 2.5 per OTU) than in fungal OTUs (0.49 ± 0.72) (Fig. 5B). In the bacterial microbiota, four patterns indicate direct microbial interactions. First, co-occurrence associations were most frequent with S2-associated OTUs, followed by S1-associated and persistent OTUs (Fig. 5C). Second, the co-occurring OTUs belonged not only to *Clostridia* and *Bacteroidia*, generally known as colonic fermenters [31], but also to non-fermenting taxa, including *Bacilli*, *Gammaproteobacteria*, and *Actinomycetia* (Fig. 5D). Third, these co-occurring OTUs showed distinct seasonal composition patterns (Fig. 5D). Fourth, negative associations (53.2 ± 0.4%) predominanted among plant glycan-degrading genera, whereas positive associations (83.0 ± 0.17%) predominated among non-plant glycan-degrading genera (Fig. 5E). These seasonally variable patterns are inconsistent with a polysaccharide-driven mechanism, further supporting season-dependent lettuce-to-gut microbial transmission and suggesting direct interactions of the transmitted microbiota with commensal gut bacteria.

**Figure 5.**
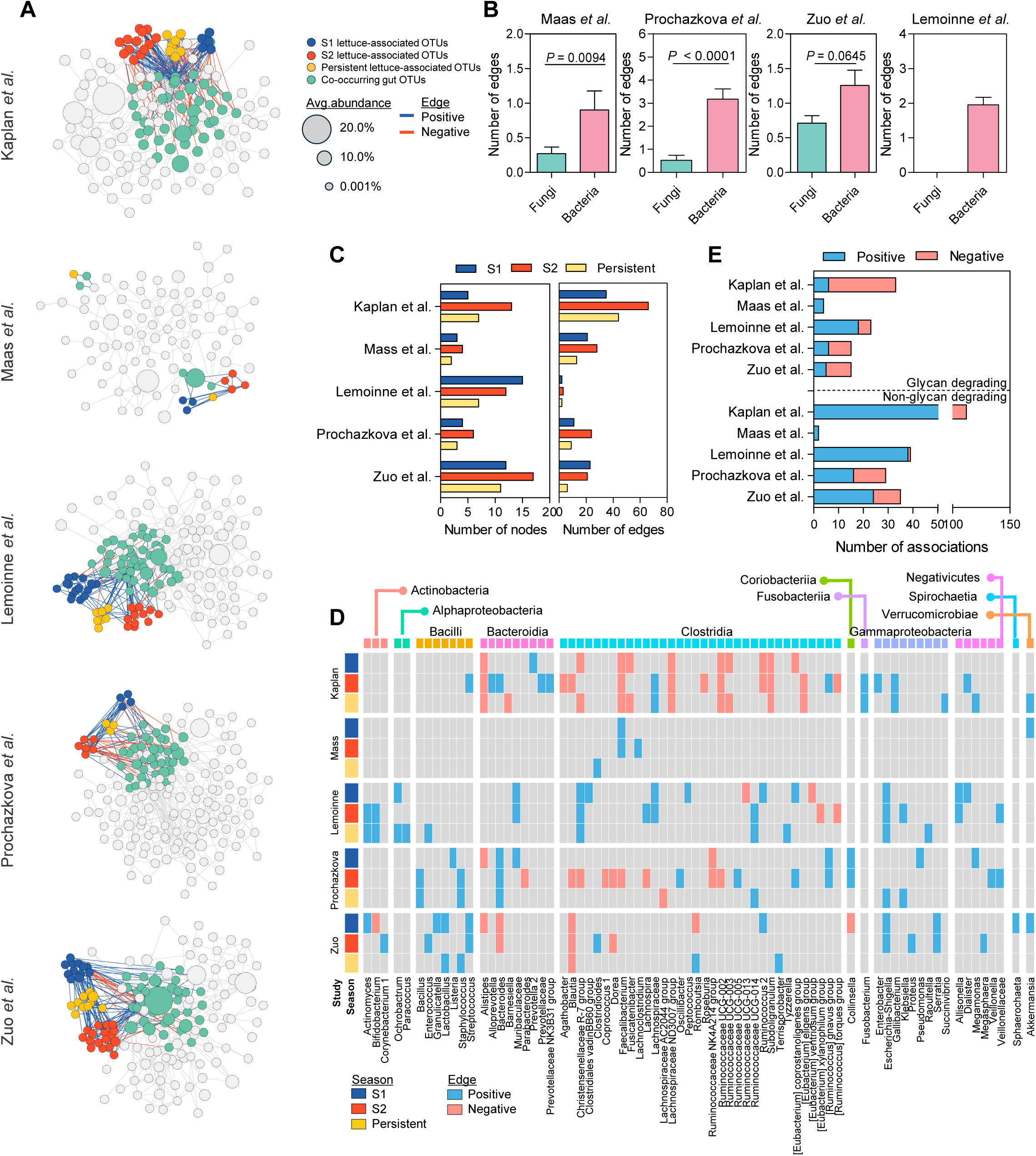
Co-occurrence associations between lettuce-associated and gut-associated OTUs. (**A)** Co-occurrence associations between lettuce-associated (S1, S2, and persistent) and gut-associated OTUs across five study cohorts. **(B)** Comparisons of edge counts between fungal and bacterial associations with gut-associated OTUs. **(C)** Node and edge counts categorized by seasonal (S1, S2) and persistent origins. **(D)** Taxonomic and seasonal profiles of co-occurring gut-associated OTUs. **(E)** Proportions of positive and negative associations categorized by glycan-degrading potential. Data are presented as mean ± s.d. Statistical significance was determined by one-tailed Mann-Whitney *U* test.

Despite these seasonal patterns, the prevalence of the lettuce-associated OTUs did not positively correlate with their prevalence in the original lettuce samples (Fig. S9C). Similar patterns were observed for OTU abundance, although partial significance was detected for the S2-specific OTUs (Fig. S9D), indicating that microbial transfer may not be strictly dose-dependent. The Sloan neutral model revealed that most lettuce-associated OTUs fell within the range of neutral expectations (89.0 ± 4.7% of fungal, 75.6 ± 5.3% of bacterial OTUs); however, a substantial proportion of bacterial OTUs deviated from neutral expectations (24.4 ± 5.3%), compared with fungal OTUs (11.0 ± 11.5%) (Fig. 4D). For example, study-shared bacterial OTUs of *Enterococcus*, *Lactococcus*, *Streptococcus*, *Clostridium sensu stricto 1*, and *Lachnoclostridium* were positively selected (Fig. 4F), whereas study-shared fungal OTUs of *Rhodotorula* and *Wallemia* appeared to follow neutral expectations (Fig. 4E). Collectively, these results suggest that lettuce-to-gut microbial transmission may be season-dependent and governed largely by neutral assembly for fungi but by both neutral and deterministic processes for bacteria.

### Seasonally variable transmission of lettuce-associated microbiota is associated with increased gut microbial diversity

We lastly asked whether the lettuce-associated microbiota is associated with changes in the gut microbiota. We compared the composition and diversity of the gut microbiota between samples containing (lettuce group) and lacking (non-lettuce group) lettuce-associated OTUs, after excluding all lettuce-associated OTUs from the OTU table and balancing group sizes through random subsampling. In most studies, bacterial composition differed significantly between the two groups, with microbial diversity significantly greater in the lettuce groups (*P* < 0.05) (Fig. 6A). A subset of the co-occurring OTUs (63.7 ± 8.2 OTUs per study) were identified as discriminant OTUs, almost all of which were enriched in the lettuce group (Fig. 6B). Accordingly, the total number of these co-occurring OTUs positively correlated with gut microbiota richness (*P* < 0.05) (Fig. S10B). Such patterns were not consistently observed in fungal composition (Fig. S11), likely because of the low co-occurrence association frequency in fungal OTUs (Fig. 5B). Taken together, these findings suggest that the seasonally variable transmission of lettuce-associated microbiota may contribute, at least in part, to shaping the composition and diversity of the human gut microbiota.

**Figure 6.**
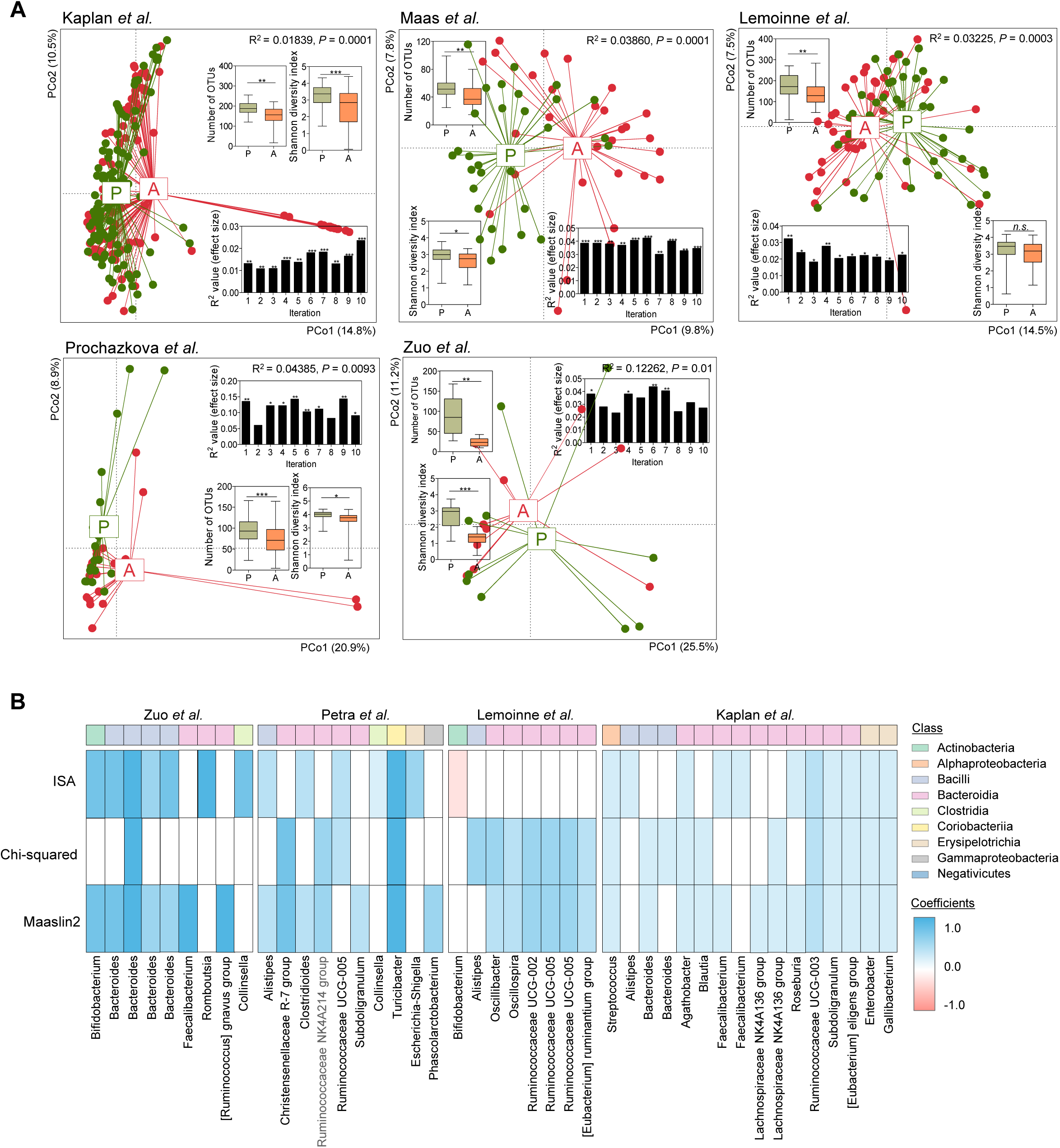
Presence of lettuce-associated microbiota shapes the composition and diversity of the human gut microbiota. **(A)** Compositional comparison of gut microbiota between individuals with and without detectable lettuce-associated bacterial OTUs. Inset plots show Shannon diversity, OTU number, and effect size for each iteration. **(B)** Heatmap of co-occurring gut OTUs identified as discriminant features between presence (P) and absence (A) groups of lettuce-associated bacterial microbiota. Data are presented as mean ± s.d. Statistical significance was determined by two-tailed Mann-Whitney *U* test and PERMANOVA. Symbol: n.s., not significant, *, *P* < 0.05; **, *P* < 0.01; and ***, *P* < 0.001.

## Discussion

Fresh vegetables serve not only as nutritional resources rich in dietary fibers, but also as microbial vectors [6, 32]. Lettuce, one of the most widely produced vegetables worldwide [33], has been documented in epidemiological studies as a vehicle for foodborne pathogens causing intestinal illness, such as *E*. *coli* O157 and *Salmonella* strains [34]. While research on the food-gut axis has traditionally focused on foodborne pathogens, attention is expanding to the commensal microorganisms associated with vegetable plants [35]. This study advances our understanding of these “edible” microbiomes in three ways [36]. First, consuming one portion of a typical lettuce salad introduces approximately 4.3 billion bacteria and 8.5 billion fungi into the gastrointestinal tract, the vast majority of which are non-pathogenic. Second, the lettuce phyllosphere microbiota exhibits a seasonal assembly pattern, leading to temporally variable microbial exposure through dietary intake. Third, lettuce-associated microbiota reaching the lower gut may interact with and enrich resident gut bacteria, particularly non-plant glycan-degraders, thereby increasing total microbiota diversity.

Although fresh vegetables harbor distinct microbiota under preharvest and postharvest conditions [37], this study focused on postharvest samples for three strategic reasons. First, the microbial diversity and ecology of postharvest lettuce microbiota remain poorly characterized [38]. Second, postharvest sampling reflects typical consumer exposure, as vegetables undergo harvesting, commercial packaging, transportation, and storage before purchase [10, 11]. Third, this approach aligned with our objective to investigate potential microbial translocation from lettuce to the human intestine. Nevertheless, seasonal variations prominently shaped both the bacterial and fungal microbiota in the postharvest lettuce phyllosphere, resulting in seasonal cycling patterns. While such patterns are expected in field-grown preharvest samples [39, 40], their persistence in postharvest samples indicates the conservation of preharvest microbiota throughout postharvest processing, likely facilitated by the short crop production cycle of lettuce [23]. Our month-based longitudinal sampling was able to capture these patterns, in contrast to the cross-sectional approaches used in previous studies [12, 41], suggesting that longitudinal analysis needs to be extended to diverse annual vegetables with varying production cycles to validate microbiota conservation between the preharvest and postharvest stages.

Seasonal cycling in the lettuce phyllosphere microbiota was driven by the interplay of stochastic and deterministic assembly processes. Stochastic processes dominated overall microbial dynamics across seasons, whereas deterministic processes favoring positive selection increasingly shaped late-season communities. Season-specific members, whose diversity increased during the late season, exhibited predominantly stochastic assembly patterns consistent with allochthonous colonization from external sources [13]. Conversely, season-persistent members, such as *Sphingomonaceae* and *Enterobacteriaceae* commonly reported in lettuce microbiota studies [42–44], displayed deterministic patterns, which are typical of core taxa in host-associated microbiomes [45], further supported by their higher prevalence and abundance compared with season-specific members (Fig. S12). These persistent members showed distinct seasonal trajectories: overall enrichment of the fungal microbiota, and selective enrichment of core taxa of the bacterial microbiota [41, 46]. The increasing diversity of season-specific members, combined with the dominance of persistent members, strengthened microbial network complexity, particularly through inter-kingdom associations [47], which may have contributed to the suppression of potentially pathogenic fungal genera in late-season microbiota, although the underlying mechanisms require further experimental investigation. Collectively, these dynamics result in seasonally increasing fungal diversity and seasonally variable bacterial compositions reaching consumers through lettuce consumption.

Fresh vegetable consumption has been associated with enhanced human health through modulation of microbial diversity and activity [48, 49], leading to the hypothesis that vegetable-associated microbiota may directly contribute to these benefits. Using multiple validation approaches, including species-level sequence homology, taxonomic consistency, phylogenetic relatedness, and environmental source tracking, this study detected lettuce-associated bacteria and fungi in the gut microbiota of approximately two-thirds and one-third of the human subjects, respectively. Twenty-four bacterial OTUs and eight fungal OTUs were shared across multiple independent studies. These detection rates may increase as additional samples from diverse cultivation regions are analyzed. These findings corroborate those of Wicaksono *et al*. [6], who demonstrated that fruit- and vegetable-derived bacteria are present in the human gut microbiome, and further extend those of David *et al*. [30] by providing evidence for the transmission of lettuce-associated fungi to the human gut. Together with our fresh vegetable virome research [16], these results suggest that bacteria, fungi, and viruses may all transfer from fresh vegetables to the human gut. While research on microbial transfer through the food-gut axis has largely focused on fermented foods [27–29], our findings support the expansion of this framework to non-fermented foods as sources of the human gut microbiota.

Lettuce-associated microbiota detected in the human gut microbiota included more season-specific members than persistent members. Furthermore, both groups of lettuce-derived bacteria co-occurred with distinct gut bacteria, predominantly with non-plant glycan-degraders. These microbial associations contributed to community-level differences between the lettuce and non-lettuce groups, characterized by increased microbial diversity through enrichment of co-abundant gut bacteria. This pattern suggests that lettuce-derived bacteria may influence the gut microbiota composition not only through simple microbial addition but also through direct interactions with resident gut bacteria [6, 50]. Given that probiotic effects can arise from low abundance [51] and the ability to transiently colonize foodborne probiotics [52], lettuce-derived bacteria could have comparable effects on the gut ecosystem. Indeed, probiotic genera such as *Lactobacillus*, *Lactococcus*, *Bacillus*, and *Enterococcus* were identified among the lettuce-associated OTUs [53], suggesting that positive health outcomes associated with dietary consumption of live microbes may extend to vegetable-associated microbiota [54, 55]. However, not all lettuce-associated bacteria and fungi reach the gut equally. The prevalence and abundance of lettuce-associated OTUs on lettuce did not correlate with those in the human gut, consistent with our previous research [16] and the findings of Mantegazza *et al*. [56] indicating that microbial translocation is not dose-dependent but selectively restricted during gastrointestinal transit. The gastrointestinal tract imposes selective barriers, such as acidic gastric fluid [57], duodenal bile acids [58], and oxygen gradients [59], that collectively require specific resistance mechanisms for microbes to reach the colon in viable form. Several lettuce-associated genera detected in the gut microbiota and showing co-occurrence associations with resident bacteria possess traits that may facilitate survival: *Acidiphilium* and *Sphingomonas* are known for acid tolerance [60]; *Bacillus* species produce resistant dormant spores [61]; members of *Bacteroidota* possess broad arrays of carbohydrate-active enzymes [62]; and *Lactococcus* spp. produce exopolysaccharides that confer protection against gastric and bile acid stress [63]. Despite these mechanistic speculations, the precise ecological processes governing this microbial transmission remain largely unresolved; understanding these processes may represent a critical step towards establishing the health benefits of vegetable-associated microbiota. In contrast, lettuce-derived fungi showed no such associations with the gut microbiota. Given that fungi are substantially less abundant and diverse than bacteria in the healthy gut [64] and that their abundance has been reported to be negatively correlated with bacterial diversity in intestinal diseases [65, 66], lettuce-derived fungi may lack the ecological capacity to establish interactions with resident gut bacteria.

Based on our findings, it can be hypothesized that fresh vegetable-associated microbiota contribute to human gut microbiome seasonality through dietary microbiota transfer. For example, seasonal gut microbiome cycling has been demonstrated in the Hadza hunter-gatherers because of seasonally variable food intake [1], and in Indian agrarian populations linked to the consumption of season-specific fermented foods [67]. Our results suggest that seasonally variable vegetable microorganisms may represent an additional contributing factor. This mechanism may be particularly relevant for industrialized populations whose diets are dominated by highly processed food [68]. For such populations, regular consumption of diverse fresh vegetables could introduce seasonally variable microbiota that partially restore the seasonal gut microbiome patterns observed in traditional rural communities, a possibility that may contribute to defining healthy gut microbiomes [2, 69, 70]. Given that each vegetable species has a unique microbiota (Fig. S2B), microbial diversity of fresh vegetables would result in the delivery of an even broader range of plant microorganisms to the gut, further amplifying such seasonal effects.

This study highlights the lettuce phyllosphere as an underappreciated source of environmental microbiota, demonstrating that dietary lettuce acts as a vector for transmitting plant-associated bacteria and fungi to the human gut. Furthermore, it provides both a theoretical framework and proof-of-concept evidence that vegetable-associated microorganisms can interact with commensal gut bacteria, potentially modulating gut microbial diversity and community structure. Our *de novo* lettuce microbiota dataset represents only a small fraction of the microbiota diversity present on fresh vegetables, and expanding microbiota databases to include a wider variety of fresh vegetables could improve the detection and tracking of vegetable-derived microbiota in the human gut. Additionally, dietary intervention experiments employing strain-level comparisons between fresh vegetable and human gut microbiomes are necessary to validate our findings and elucidate the extent and mechanisms underlying these microbial transfers. Overall, this study advances our understanding of fresh vegetable microbial ecology and its relationships with the human gut microbiota. By integrating the edible plant microbiome into the “One Health Microbiome” [71], these findings provide a foundation for future investigations into the impact of food-associated microbiota on human health through the food-gut axis.

## Acknowledgments

We sincerely thank Prof. Koen Venema, Ruixue Dai, and the CHILD Cohort Study for providing the gut microbiome metadata.

## Author contributions

S.A.I., E.J.P. and M.S.K. conceived and designed the experiments; S.A.I, J.W.P., Y.E.Y. and Y.E.L performed the experiments; S.A.I. and M.S.K. analyzed the data; S.A.I, J.W.P., E.J.P., and M.S.K. contributed materials/analysis tools; S.A.I., E.J.P. and M.S.K. wrote the manuscript.

## Data availability

The EMBL-EBI accession numbers for the sequences of the ITS2 region, the V5-V6 regions, and the V3-V4 regions of the 16S rRNA gene are PRJEB96271, PRJEB96269, and PRJEB96270, respectively.

