## Supplementary methods and figures for "Seasonal dynamics in lettuce phyllosphere microbiota and potential transmission to the human gut"

**This PDF file includes:**

Supplementary Materials and Methods

Supplementary Figures 1 to 12

### Supplementary Materials and Methods

*Quantification of viable cells of fungi and bacteria*

Total numbers of viable fungal and bacterial cells were determined by counting colony-forming units (CFUs). Each sample mixture was ten-fold serially diluted and spread on tryptic soy agar (TSA, BD, USA) for analysis of total bacteria and potato dextrose agar (PDA, BD, USA) for analysis of total fungi. PDA was supplemented with chloramphenicol (50mg/L; Sigma-Aldrich, Germany) to inhibit bacterial growth [1] and Rose Bengal (Sigma Aldrich, Germany) to suppress fungal hyphal formation [2]. Plates were incubated at 30 °C for five days for fungi and three days for bacteria.

*PCR conditions*

PCR was conducted using AccuPower PCR PreMix (Bioneer, Korea) under the following conditions: initial denaturation at 95 °C for 2 min; 35 cycles at 95 °C for 30 sec, 56 °C for 30 sec, and 72 °C for 30 sec; and a final extension at 72 °C for 10 min for ITS2 region. 94 °C for 3 min; 28 cycles at 94 °C for 30 sec, 55 °C for 30 sec, and 72 °C for 30 sec; and a final extension step at 72 °C for 5 min for V5-V6 regions of 16S rRNA. 94 °C for 3 min; 28 cycles at 94 °C for 30 sec, 50 °C for 30 sec, and 72 °C for 30 sec; and a final extension step at 72 °C for 5 min for V3-V4 regions of 16S rRNA. ITS2 and V5-V6 amplicons are combined from five replicates, and V3-V4 are from three replicates using QIAquick PCR purification Kit (Qiagen, Germany).

*Amplicon sequencing and sequence analysis*

The internal transcribed spacer region 2 (ITS2) of the ribosomal RNA gene was amplified using fITS7 and ITS4 primers [3] with Illumina sequencing overhang adaptors, as described previously [4]. The V5–V6 regions of the 16S rRNA gene were amplified using chloroplast-excluding 799f and 1115r primers [5]. Additionally, the V3-V4 regions of the 16S rRNA gene were amplified using 341f and 805r primers, as described previously [6]. Sequencing was performed at Macrogen (Korea) for ITS2 and V5-V6 amplicons, and at Novogene (China) for V3-V4 amplicons. Taxonomy was assigned using a Naïve Bayes classifier trained on the UNITE (version 8.3, release 2021-05-10) and SILVA (version 138.1, release 2020-11-02) databases, both at the 99% threshold. ASVs assigned to unclassified taxa, mitochondria or chloroplasts were removed. Samples were rarefied to 20,000 reads for fungi and 3,000 reads for bacteria, and ASVs with less than 0.05% relative abundance were removed. The observed ASVs, Shannon index, and Pielou's evenness were estimated for alpha-diversity. The log-transformed Bray-Curtis dissimilarity and Jaccard distance were calculated for beta-diversity. For a global comparison of microbiomes in fresh fruits and vegetables, the ITS and 16S rRNA gene sequence datasets were retrieved from previous studies [7, 8]. The downloaded datasets were analyzed as described above.

*Quantification of internal transcribed spacer 2 region and 16S rRNA gene sequences*

The gene copy number of bacterial 16S rRNA and fungal ITS2 region was quantified with primer pairs specific for the V5-V6 regions of 16S rRNA gene [5] and ITS2 region [9] using SYBR Green Supermix (Bio-Rad, Hercules, CA). Genomic DNAs of *Escherichia coli* ER2925 (2.5 ng/μl) and *Saccharomyces cerevisiae* BY4741 (1.0 ng/μl) were quantified with the same primer pairs to make standard curves. All reactions were performed in triplicate.

*Normalization of gene copy numbers*

Measured gene copy numbers were normalized using the average rRNA gene copy numbers of most abundant phyla: 5.8 copies for *Gammaproteobacteria* [10] and 4.7 and 7.2 copies for *Ascomycota* and *Basidiomycota*, respectively [11].

*Meteorological parameters*

The monthly average meteorological values of the precipitation rate (mm/day), temperature (°C), relative humidity (%), wind speed (m/s), solar radiation ($W/m^2$), and soil wetness (wfv) were obtained from the NASA Prediction of Worldwide Energy Resources (https://power.larc.nasa.gov)on the basis of the harvest cities and sample collection dates indicated on the sample packaging **(Table S8)**.

*Logistic regression for season-associated ASV detection*

Logistic to presence-absence ASV profiles were generated for persistent microbiota. Abundances were first binarized (1 = present, 0 = absent), and for each ASV, a logistic model was fitted using the '*glm*' function in R, with season as the explanatory variable. ASVs with season-associated coefficients showing *P*-value < 0.05 were considered significantly skewed. The number of ASVs enriched in each season was summarized, and seasonal skewness was visualized using density plots of log-transformed presence ratios.

*Co-occurrence network analysis*

Single-kingdom and cross-kingdom microbial co-occurrence networks were constructed using SPIEC-EASI [12]. ASVs with less than 0.1% average abundance within each seasonal cluster were filtered out. The Meinshausen-Buchmann neighborhood selection method was selected based on the Stability Approach to Regularization Selection (StARS). The StARS variability threshold was set to 0.01 for all networks with 99 repetitions and a lambda of 20 to achieve network stability. The number of nodes and edges, average number of neighbours and characteristic path length were calculated using the NetworkAnalyzer plugin of Cytoscape (version 3.10.2). the co-occurrence networks of lettuce- and gut-associated OTUs, gut-associated OTUs with less than 0.1% average abundance were filtered out and combined with lettuce-associated OTUs.

*Sloan Neutral Model*

The Sloan Neutral Model (SNM) was used to evaluate the contribution of neutral processes in microbial community assembly [13]. The SNM was constructed using '*pctax*' package in R based on the rarefied ASV abundance table. The model estimates the immigration rate (*m*) and evaluates the goodness-of-fit ($R^2$) to indicate how well neutral dynamics explain observed community structures.

*Potential pathogen and food spoiler identification*

Potential fungal pathogens were identified at the genus level based on Funguilds (Animal pathogen, Plant pathogen) [14], FungalTraits (Plant pathogen, Animal parasite, Lichen parasite, Mycoparasite, Algal parasite, and Arthropod parasite) [15], and ResNets (Human pathogen, Animal pathogen, Plant pathogen) [16]. Potential bacterial pathogens were assigned at the genus level based on MBPD (Plant pathogens, Animal pathogens, Zoonotic pathogens) [17]. Spoilage fungi and bacteria associated with fresh fruits and vegetables were identified at the genus level, as described in previous studies [18-32].

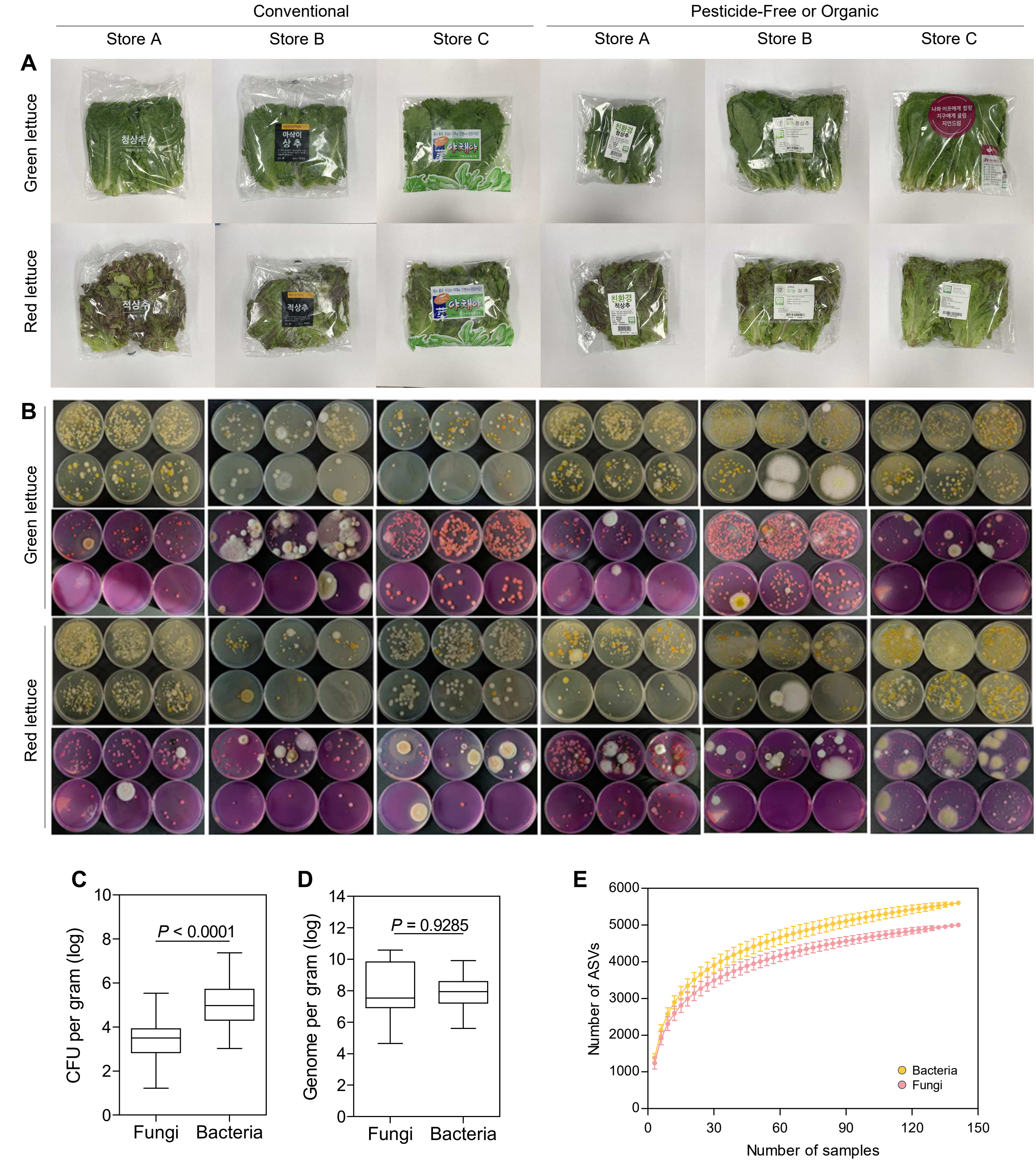


**Supplementary Figure 1. Characterization of lettuce-associated microbiota. (A)** Representative images of green (above line) and red (below line) lettuce samples collected from six different grocery stores in Daejeon, Korea. **(B)** Viable fungal and bacterial colonies cultured from lettuce samples: bacterial colonies grown on TSA plates (top line) and fungal colonies on PDA plates (bottom line), shown at dilution factors of $10^{-4}$ and $10^{-5}$ for bacteria and $10^{-2}$ and $10^{-3}$ for fungi. **(C)** Comparison of total viable fungi and bacteria cell counts. **(D)** Comparison of total fungal and bacterial genome numbers. **(E)** Species accumulation curve of fungal and bacterial ASVs. All data are mean ±s.d. Statistical significance was determined by two-tailed Mann-Whitney tests.

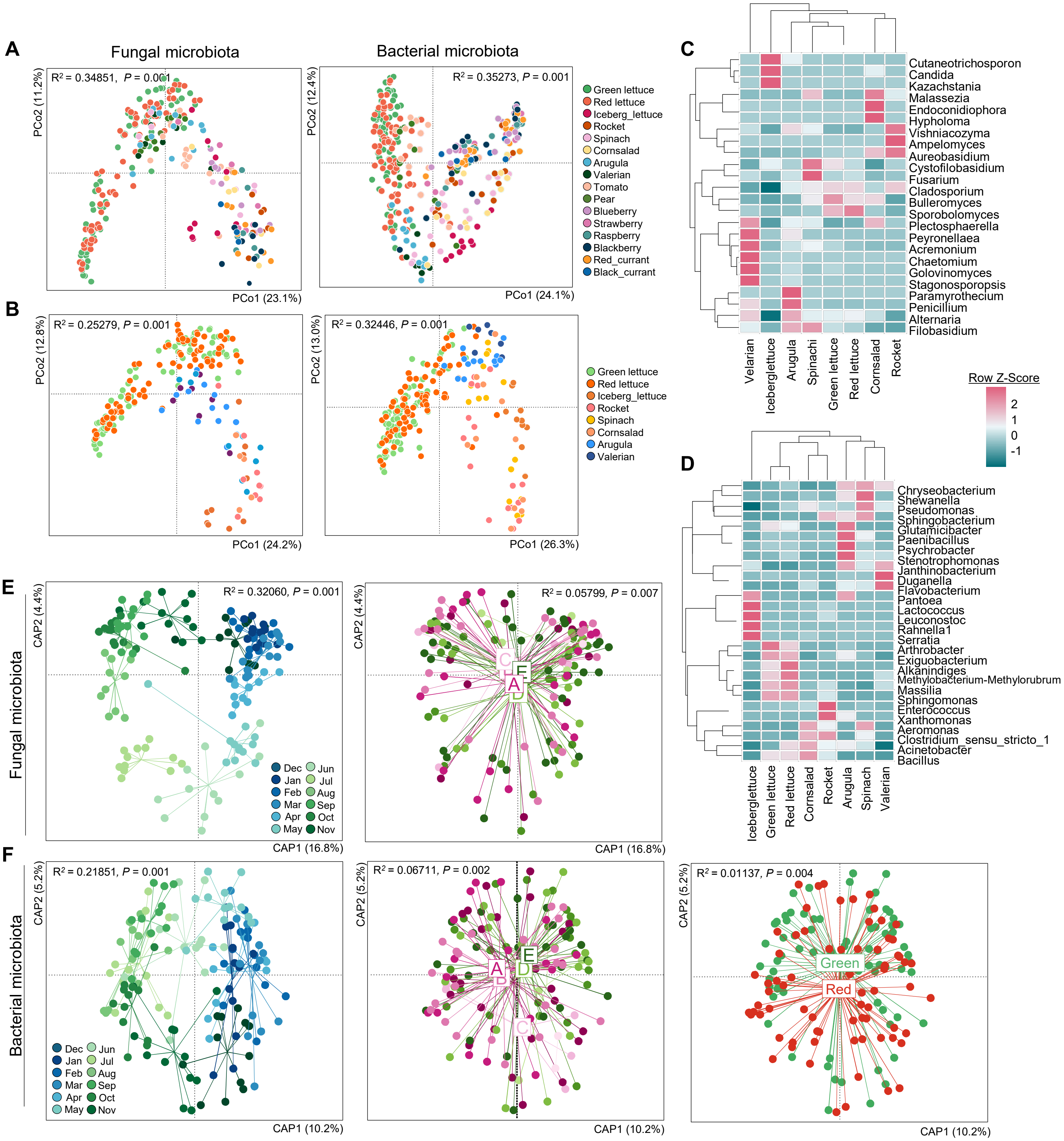


**Supplementary Figure 2. Compositional structure and environmental drivers of lettuce-associated microbiota. (A)** Global comparison of fungal (left panel) and bacterial (right panel) microbiota with those on other fruits and vegetables, and **(B)** with only fresh vegetables. **(C)** Heatmaps of discriminant fungal and **(D)** bacterial genera distinguishing from other leaf vegetable types shown in **(B)**. **(E, F)** Redundancy analysis of fungal microbiota based on sampling month (left panel) and distributor (right panel) and of bacterial microbiota based on sampling month (first panel), distributor (second panel), and host species (third panel). Redundancy analysis was performed based on log-transformed Bray-Curtis dissimilarity matrices. Statistical significance was determined by PERMANOVA.

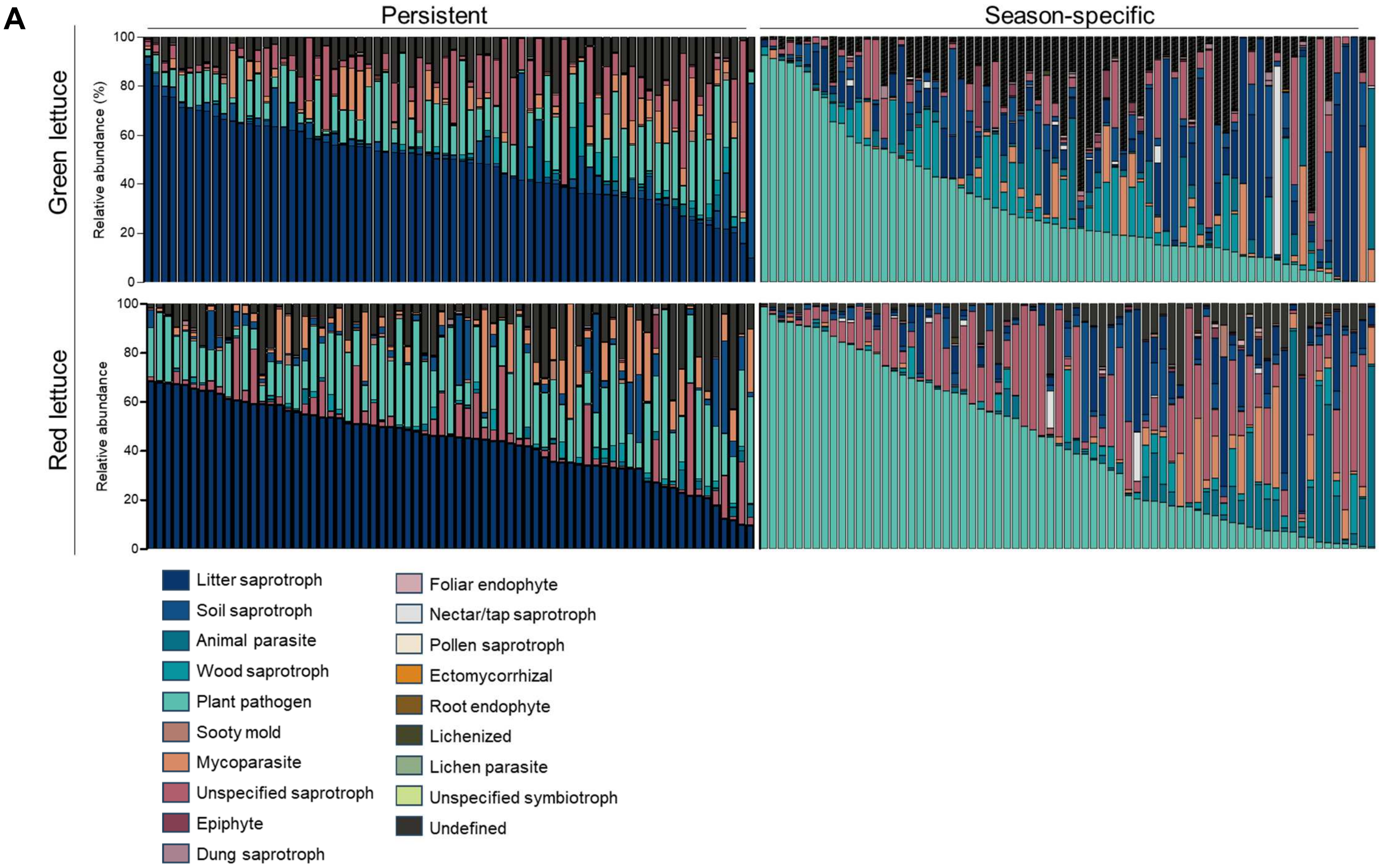


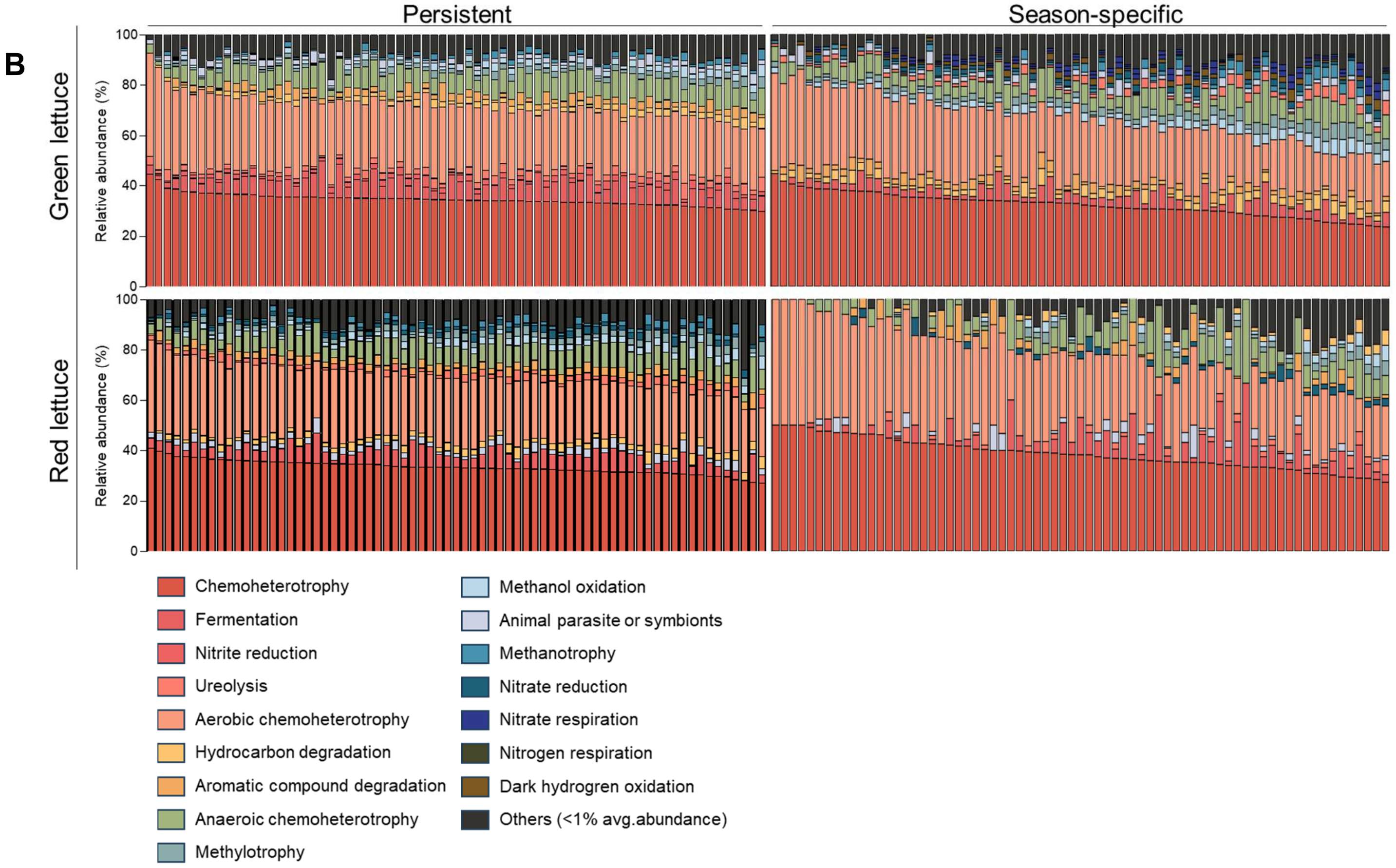


**Supplementary Figure 3.** Functional guild profiles of and season-specific and persistent taxa in **(A)** fungal and **(B)** bacterial microbiota, assigned by FungalTraits and FAPROTAX, respectively.

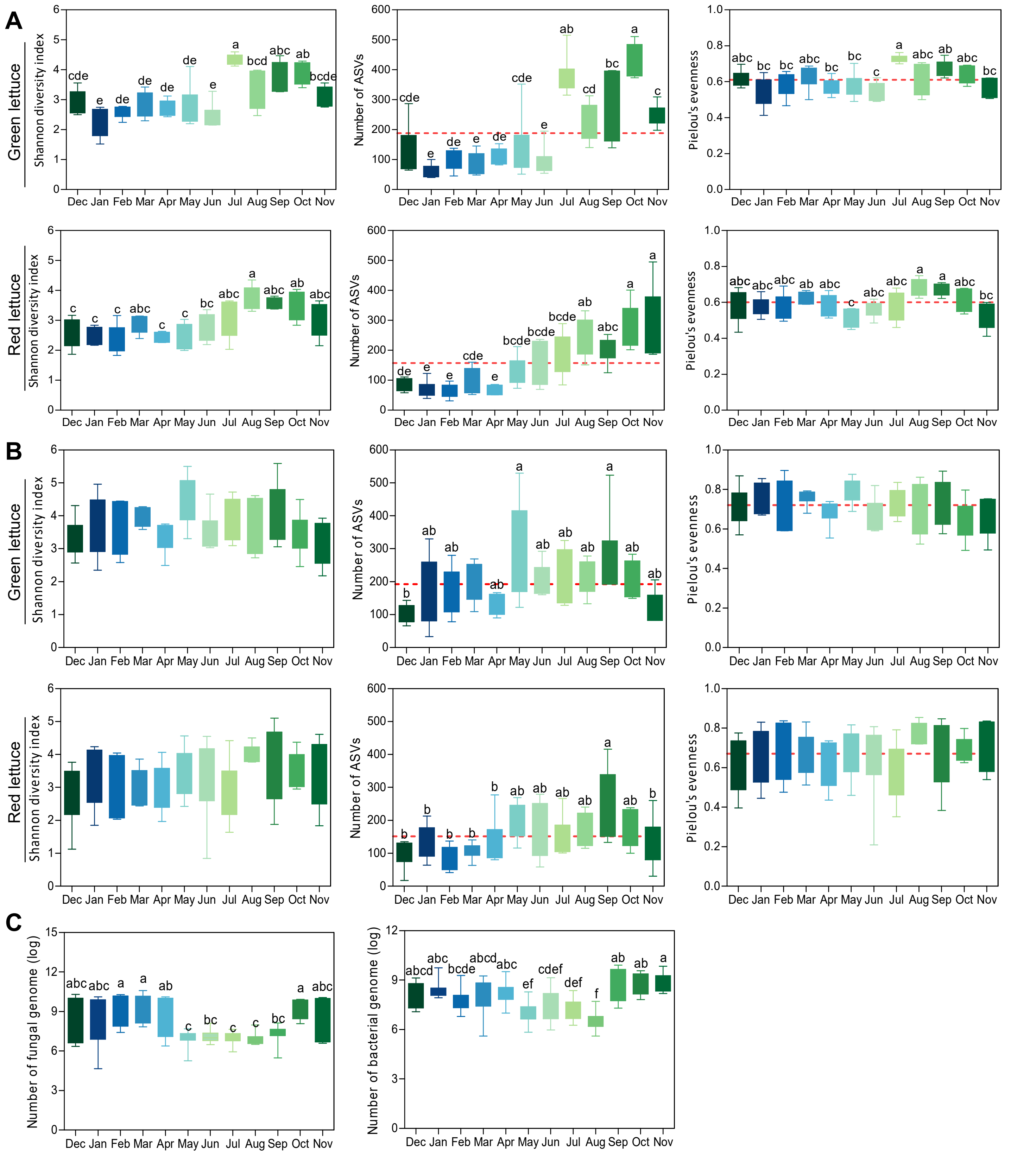


**Supplementary Figure 4. Microbial diversity and total abundance of lettuce-associated fungal and bacterial microbiota. (A)** Microbial diversity of fungal and **(B)** bacterial microbiota according to sampling month. **(C)** Comparison of genome numbers for fungi (left panel) and bacteria (right panel). All data are presented mean ± s.d. Statistical significance was determined by one-way ANOVA with Tukey's multiple comparison test.

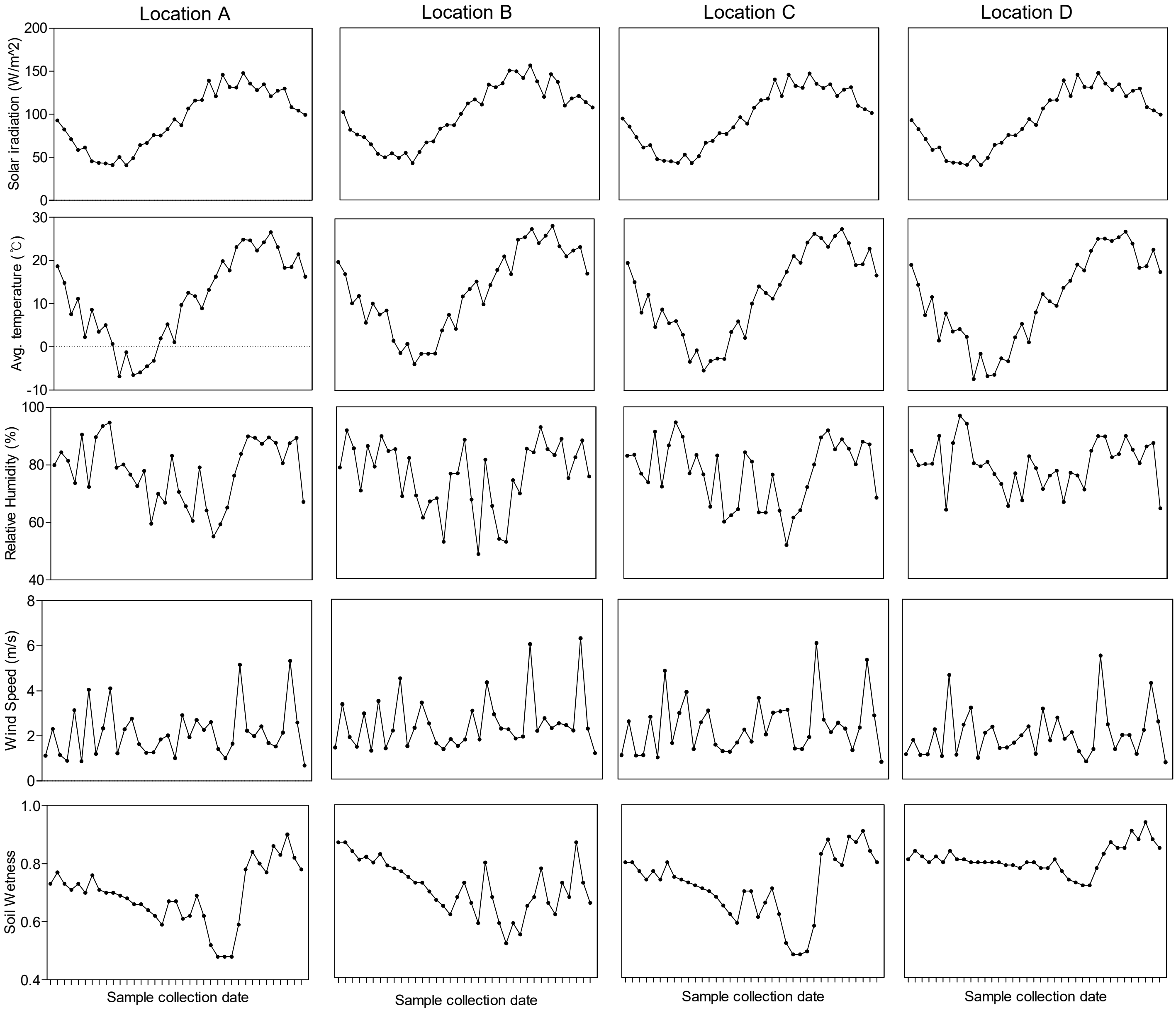


**Supplementary Figure 5. Temporal variation in agro-meteorological factors across lettuce-harvesting locations.** Temporal variation in agro-meteorological factors (solar irradiation, average temperature, relative humidity, wind speed, soil wetness) across four lettuce-harvesting cities (A, B, C, and D). All data were obtained from the NASA Predition Of Worldwide Energy Resources.

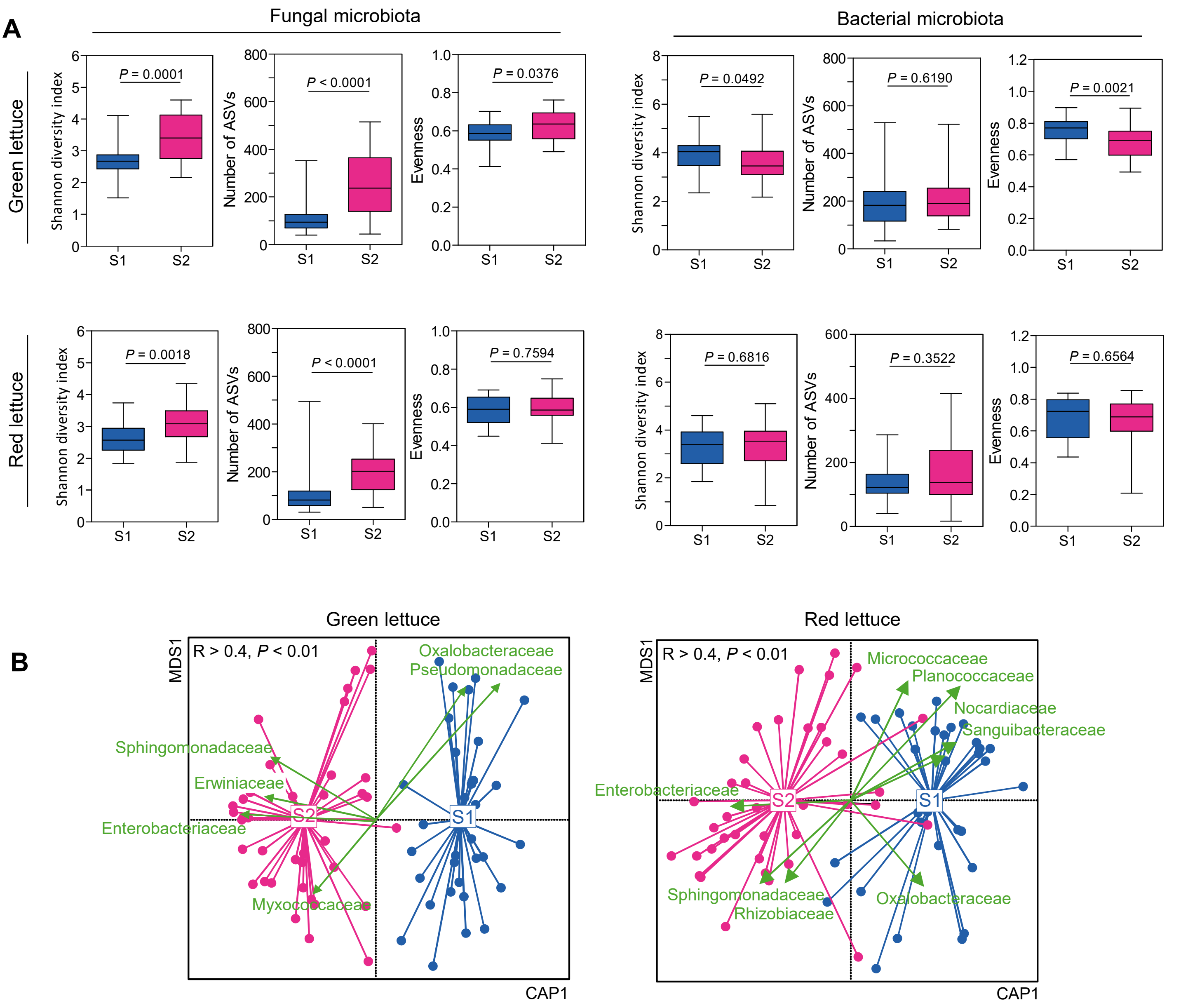


**Supplementary Figure 6. Microbial diversity and taxonomic enrichment between seasonal clusters. (A)** Microbial diversity of fungal and bacterial microbiota within each seasonal cluster. **(B)** RDA plots and discriminant bacterial families across seasons. Data are presented as mean ± s.d.. Statistical significance was determined by two-tailed Mann-Whitney U test and Spearman correlation.

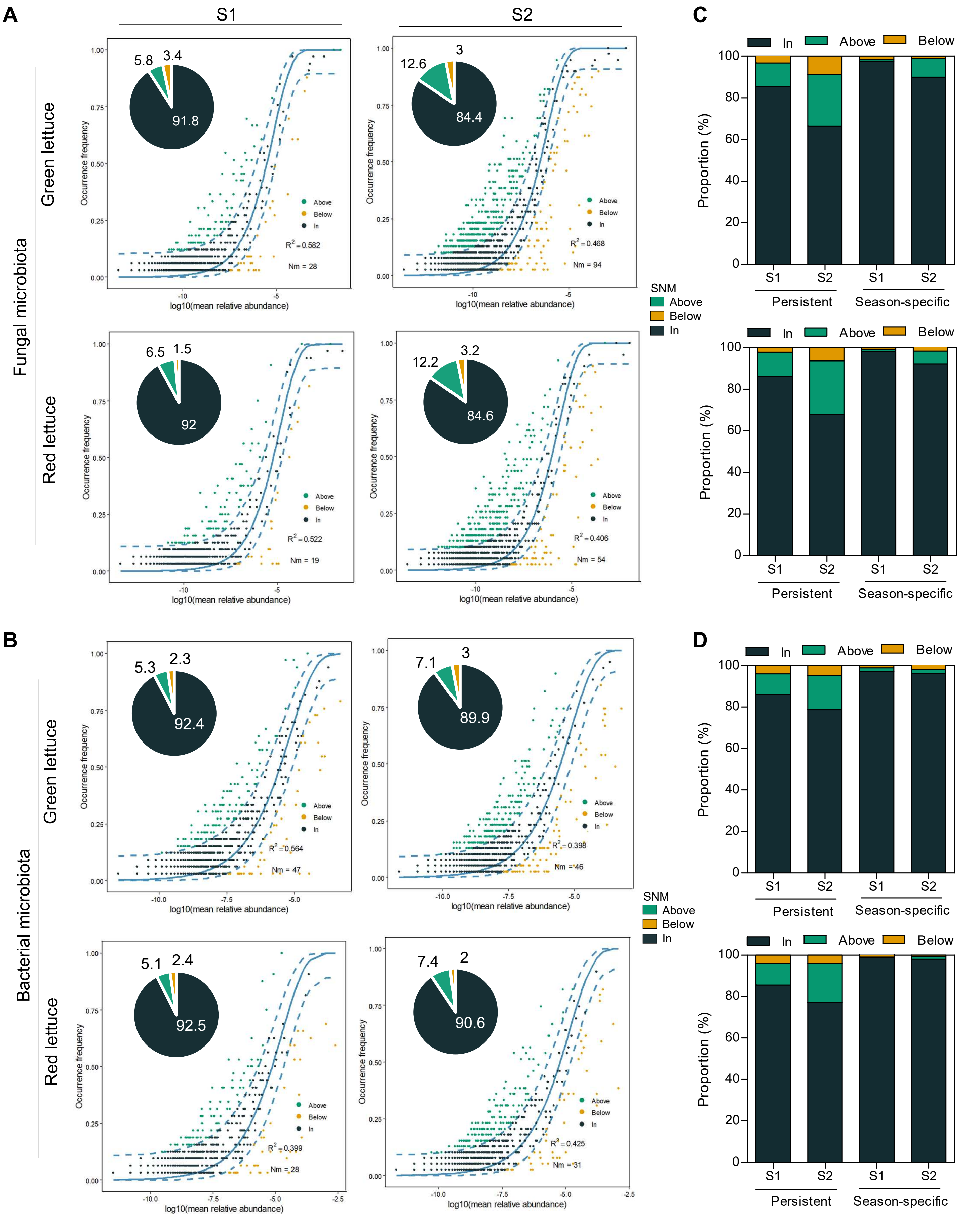


**Supplementary Figure 7. Neutral community modeling and persistence of lettuce-associated microbial ASVs.** Sloan Neutral Model (SNM) fit for **(A)** fungal and **(B)** bacterial microbiota. Pie charts show the proportions of ASVs falling within, above, or below model predictions. Proportions of persistent and season-specific ASVs categorized by SNM status for **(C)** fungal and **(D)** bacterial microbiota.

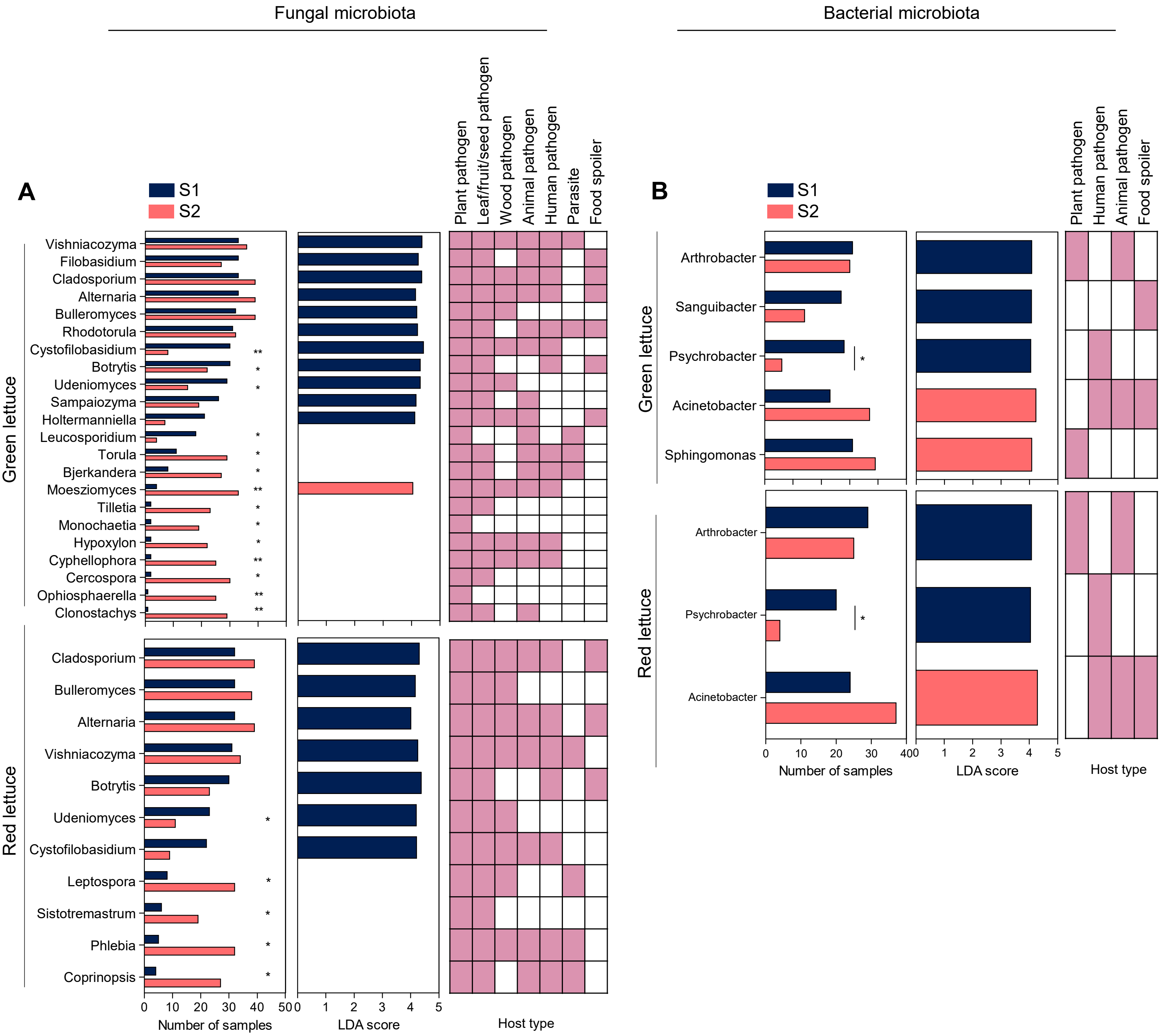


**Supplementary Figure 8. Potentially opportunistic pathogenic microbial genera in lettuce.** Potentially opportunistic pathogenic **(A)** fungal and **(B)** bacterial genera. Panels show the number of samples in which each genus was detected (left), LDA scores (middle), and host types (right). Statistical significance was determined by Chi-square test. Symbol: *, $P < 0.05$; **, $P < 0.01$.

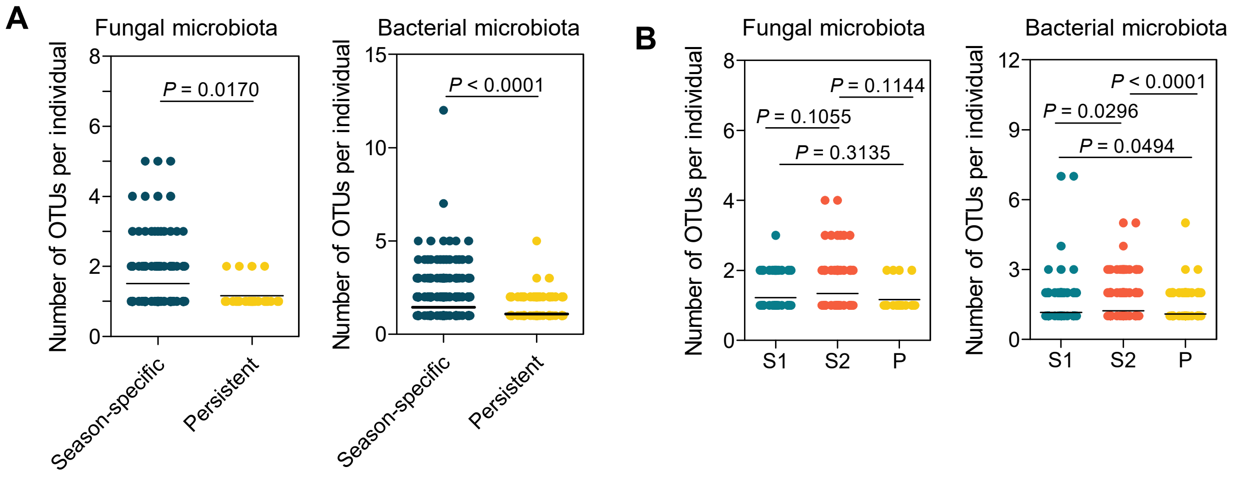


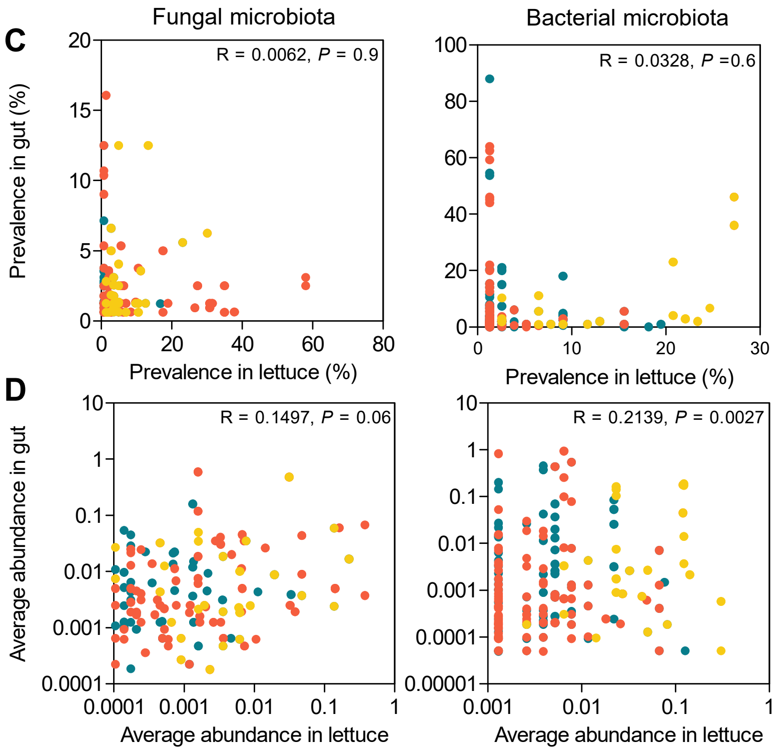


| | Fungal microbiota | Bacterial microbiota |
|---|---|---|
| S1 | R = -0.1661, *P* = 0.2 | R = -0.012, *P* = 0.3 |
| S2 | R = 0.0317, *P* = 0.7 | R = -0.057, *P* = 0.5 |
| P | R = 0.1541, *P* = 0.3 | R = 0.3182, *P* = 0.1 |

| | Fungal microbiota | Bacterial microbiota |
|---|---|---|
| S1 | R = 0.0196, *P* = 0.8 | R = 0.0901, *P* = 0.3 |
| S2 | R = 0.2485, *P* = 0.0085 | R = 0.1916, *P* = 0.04 |
| P | R = 0.0818, *P* = 0.6 | R = 0.0919, *P* = 0.6 |

**Supplementary Figure 9. Gut microbial OTU counts and correlation with lettuce microbiota at the individual level.** Number of OTUs per individual categorized by **(A)** season-specific and persistent groups and **(B)** S1, S2, and persistent groups. Spearman correlations of **(C)** prevalence and **(D)** average abundance between lettuce-associated and human gut OTUs. The table shows the Spearman correlation coefficients and *P*-values for seasonal microbiota. Data are presented as mean ± s.d. Statistical significance was determined by one-tailed Mann-Whitney *U* test and Spearman correlation.

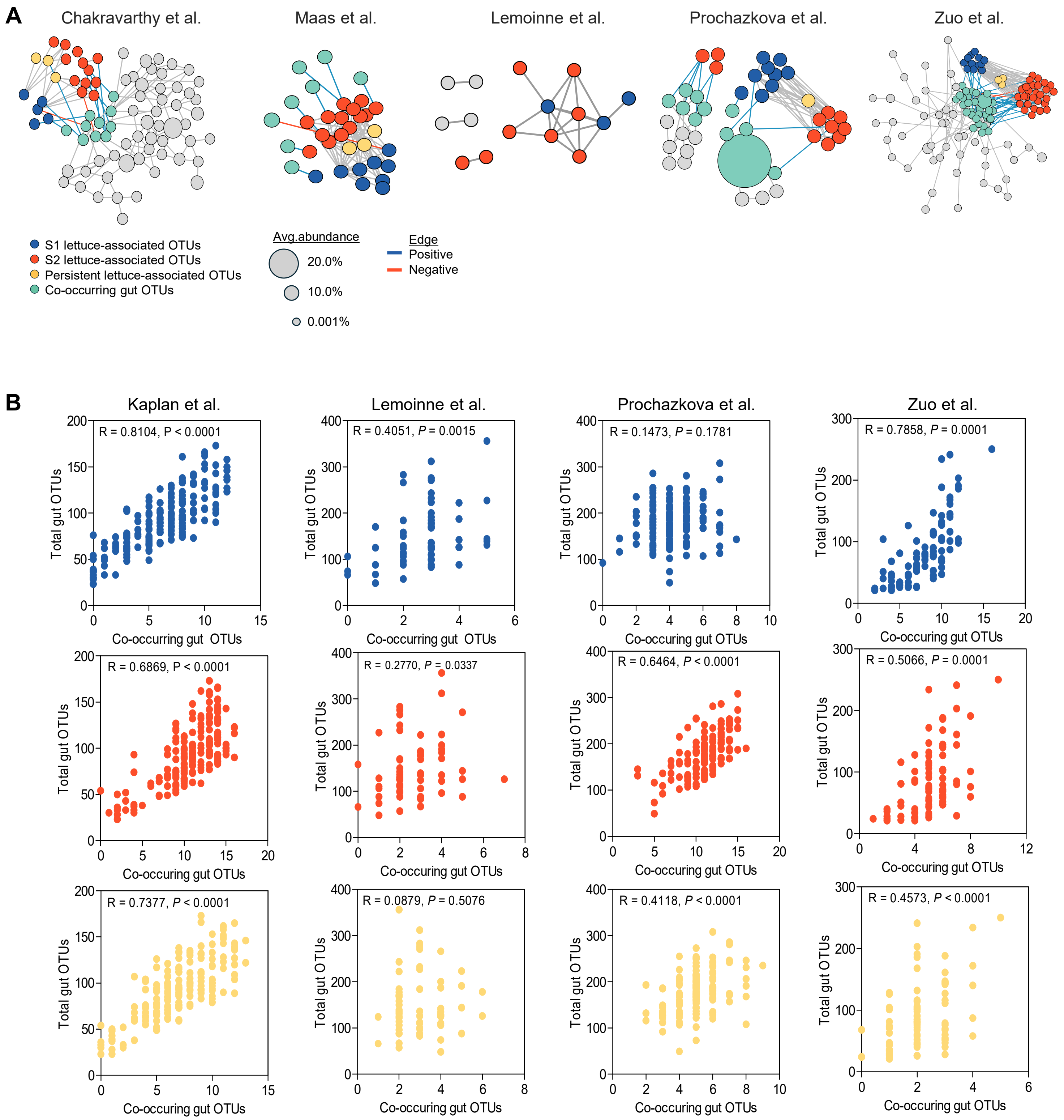


**Supplementary Figure 10. (A)** Co-occurrence networks of lettuce-associated (S1, S2, and persistent) and gut-associated fungal OTUs. **(B)** Spearman correlations between total gut bacterial OTU richness and the numbers of seasonal (S1, S2) and persistent co-occurring gut bacterial OTUs. Statistical significance was determined by Spearman correlation.

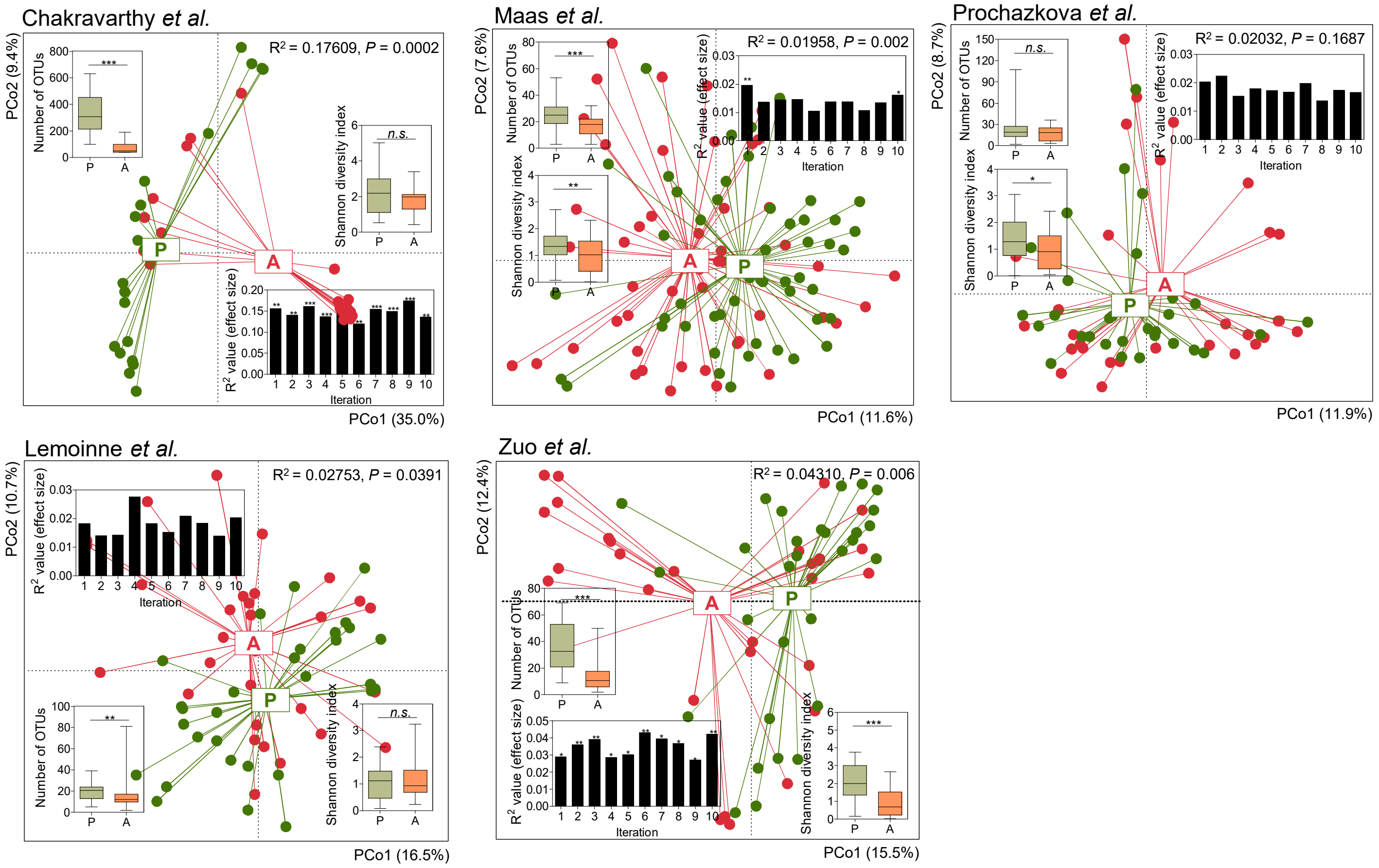


**Supplementary Figure 11. Gut fungal diversity comparisons between individuals with and without lettuce-associated fungal OTUs.** Compositional comparison of gut fungal microbiota between groups containing and not containing lettuce-associated fungal OTUs. Inset plots show Shannon diversity, OTU number, and effect size for each iteration. Data are presented as mean ± s.d. Statistical significance was determined by two-tailed Mann-Whitney *U* test and PERMANOVA. Symbol: n.s., not significant; *, $P < 0.05$; **, $P < 0.01$; and ***, $P < 0.001$.

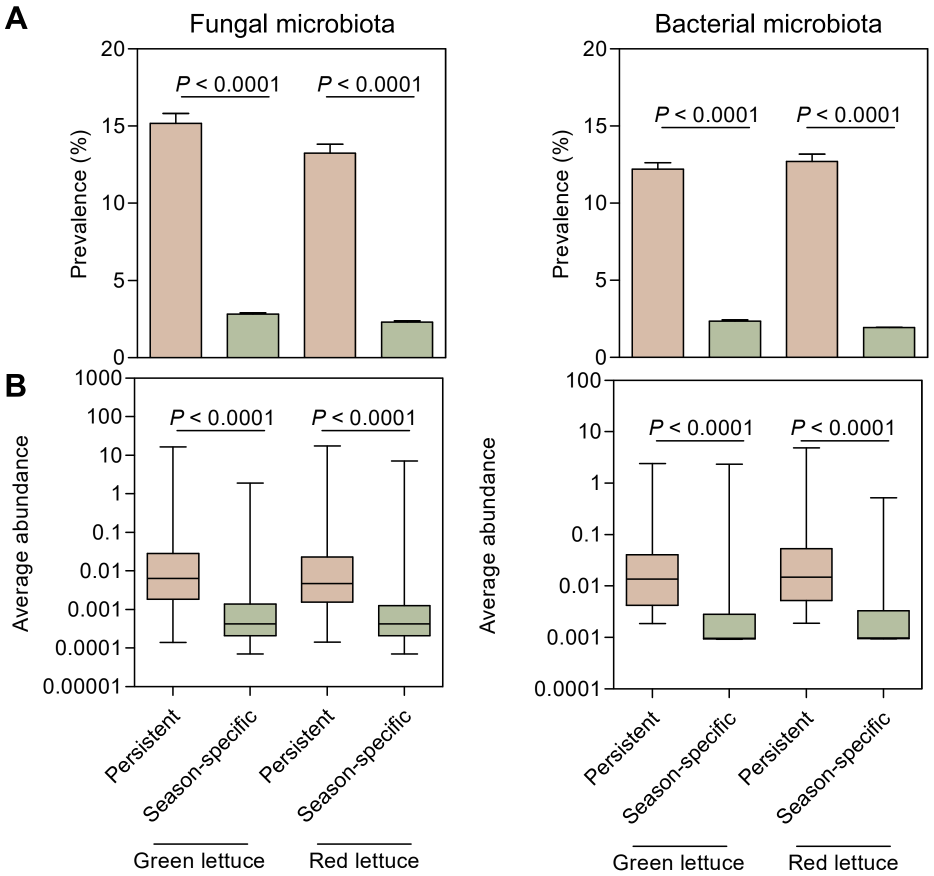


**Supplementary Figure 12. Prevalence and abundance of persistent and season-specific microbiota. (A)** Prevalence and **(B)** average abundance of fungal and bacterial microbiota compared between persistent and season-specific microbiota. Data are presented as mean ± s.d. Statistical significance was determined by two-tailed Mann-Whitney *U* test.
